# Role of the medial prefrontal cortex in the effects of rapid acting antidepressants on decision-making biases in rodents

**DOI:** 10.1101/2020.01.22.915132

**Authors:** CA Hales, JM Bartlett, R Arban, B Hengerer, ESJ Robinson

**Affiliations:** School of Physiology, Pharmacology and Neuroscience, Faculty of Biomedical Sciences, University of Bristol, Bristol, BS8 1TD, UK; CNS Diseases Research, Boehringer Ingelheim GmbH & Co. KG, Biberach an der Riss, Germany

**Author notes:** **Corresponding author:** Name: Prof. Emma Robinson, Address: School of Physiology, Pharmacology and Neuroscience, Faculty of Biomedical Sciences, Tankards Close, University of Bristol, Bristol, BS8 1TD, UK.

## Abstract

Major Depressive Disorder is a significant and costly cause of global disability. Until the discovery of the rapid acting antidepressant (RAAD) effects of ketamine, treatments were limited to drugs that have delayed clinical benefits. The mechanism of action of ketamine is currently unclear but one hypothesis is that it may involve neuropsychological effects mediated through modulation of affective biases (where cognitive processes such as learning and memory and decision-making are modified by emotional state). Previous work has shown that affective biases in a rodent decision-making task are differentially altered by ketamine, compared to conventional, delayed onset antidepressants. This study sought to further investigate these effects by comparing ketamine with other NMDA antagonists using this decision-making task. We also investigated the subtype selective GluN2B antagonist, CP-101,606 and muscarinic antagonist scopolamine which have both been shown to have RAAD effects. Both CP-101,606 and scopolamine induced similar positive biases in decision-making to ketamine, but the same effects were not seen with other NMDA antagonists. Using targeted medial prefrontal cortex (mPFC) infusions, these effects were localised to the mPFC. In contrast, the GABA_A_ agonist, muscimol, induced general disruptions to behaviour. These data suggest that ketamine and other RAADs mediate a specific effect on affective bias which involves the mPFC. Non-ketamine NMDA antagonists lacked efficacy and we also found that temporary inactivation of the mPFC did not fully recapitulate the effects of ketamine, suggesting a specific mechanism.

## Introduction

Major Depressive Disorder (MDD) is a prevalent psychiatric disorder, affecting over 300 million people globally^1^. It is the leading worldwide cause of disability, and, until recently, pharmacological treatments were limited to drugs that take weeks to improve symptoms and subjective reporting of mood^2^. The discovery of the rapid acting antidepressant (RAAD) effects of ketamine, an NMDA receptor antagonist, has rejuvenated the field by demonstrating that subjective changes in mood in depressed patients can be seen less than 2 hours following administration and are sustained for at least 7 days in some patients^3^. Although this RAAD has been shown repeatedly^4,5,6,7,8^, the mechanism underlying this effect is unclear, and better understanding could be critical for the development of new, fast-acting treatments.

Patients with MDD exhibit affective biases, whereby impairments in emotional processing leads to reduced positive and/or enhanced negative biases in multiple cognitive domains, including attention, memory, emotional interpretation and decision-making^9,10,11^. In humans, acute (and chronic) treatment with conventional antidepressants induces positive biases in emotional memory and recognition in healthy controls^12,13,14^ and patients^15^, despite a lack of subjectively reported change in mood. It has been suggested that similar affective biases can also be measured in non-human animals using learning and memory tasks^16^ and in decision making under ambiguity (first demonstrated by Harding et al.^17^ using a judgement bias task). For review and more detailed discussion of translational studies of affective biases see Robinson and Roiser^18^. Judgement bias tasks (also known as cognitive bias tasks, or ambiguous cue interpretation tasks) were first developed as a cognitive test to measure animal affect (see reviews by Mendl et al.^19^ and Roelofs et al.^20^). In the task, animals are trained to associate the presentation two distinct reference cues with two differently valenced outcomes (e,g. positive: reward/high reward, or negative/less positive: punishment/low reward). After training, individuals are presented with untrained, ambiguous cue(s), and responses to these are measured to see whether they respond with a positive or negative bias (more responses matching the positive or negative choice respectively). A recent systemic review and meta-analysis across judgement bias tasks in animals has shown that across 20 published research articles, pharmacological manipulations to induce changes in affective state overall did alter decision making about ambiguous cues as predicted^21^, demonstrating the validity of these types of tasks. In previous work in rodents in our lab using a reward-based judgement bias task (first reported by Hales et al.^22^), decision making biases were differentially altered by conventional, delayed acting antidepressants versus the RAAD ketamine^24^. In this task, where reference cues are associated with more or less positive outcomes^22–24^, we found that an acute, low dose of ketamine, but not acute treatment with another NMDA receptor antagonist, PCP, immediately induced more optimistic decision making, the direction that would be induced by a more positive affective state, whereas acute treatment with conventional antidepressants had no effect on bias^24^. However, when given chronically, the conventional antidepressant fluoxetine did induce a positive bias^24^, but only over a timescale similar to the drugs’ efficacy in patients, as measured by self-reported improvements in symptoms and mood^25^. The same pattern was also seen in this task with negative affective states, where a chronic stress manipulation, but not an acute stressor, induced more pessimistic decision making at later timepoints^22^.

The aim of this study was to build upon these findings by testing a selection of other drugs that act via NMDA receptor antagonism: lanicemine, a low-trapping NMDA receptor channel blocker developed for the treatment of MDD, but failed to show efficacy in clinical trials^26^; memantine, an Alzheimer’s medication that is a moderate affinity, non-competitive NMDA receptor antagonist, but also lacked antidepressant efficacy in clinical trials^5,27^; and MK-801, a potent, non-competitive NMDA receptor antagonist that has shown RAAD efficacy in animal models^28^. We also tested other compounds that have been shown to have RAAD in human clinical trials: the GluN2B subunit selective NMDA receptor antagonist CP-101,606^29^, and the acetylcholine muscarinic receptor antagonist scopolamine^30^. We also tested additional doses of ketamine and PCP to ensure we had examined effects across a wider range of receptor occupancy and in line with doses commonly used in preclinical animal models used to study depression^31^. To investigate the mechanism underlying the rapid positive change in decision-making bias we tested local administration of drugs shown to cause this effect directly into the prefrontal cortex (PFC), a brain area thought to be critical in the mechanism of RAAD of ketamine^32,33^ and previously shown to modulate learning biases in rodents^34^.

## Materials and Methods

### Animals and apparatus

Three cohorts of male Lister Hooded rats (each cohort n=16) were used (Envigo, UK). Rats were pair-housed with environmental enrichment, consisting of a red 3 mm Perspex house (30×10×17cm), a large cardboard tube (10cm diameter), a wood chew block (9×2.5×2.5cm) and a rope tied across the cage lid (the rope was not present in cages for cohort 3 post-surgery to avoid any possibility of implanted cannula getting caught). Animals were kept under temperature (19-23°C) and humidity (45-65%) controlled conditions on a 12-h reverse lighting cycle (lights off at 08:00h). Water was available *ad libitum* in the home cage, but rats were maintained at no less than 90% of their free-feeding body weight, matched to a standard growth curve, by restricting access to laboratory chow (LabDiet, PMI Nutrition International) to ~18g per rat per day. All procedures were carried out under local institutional guidelines (University of Bristol Animal Welfare and Ethical Review Board) and in accordance with the UK Animals (Scientific Procedures) Act 1986. Rats weighed 270-305 g (cohort 1) / 250-295 g (cohort 2) / 240-290 g (cohort 3) at the start of training, and 400-465 g (cohort 1) / 360-460 g (cohort 2) / 320-380 g (cohort 3) by the start of experimental manipulations. During experiments all efforts were made to minimise suffering including using a low stress method of drug administration^35^, and at the end of experiments rats were killed by giving an overdose of sodium pentobarbitone (200mg/kg). Behavioural testing was carried out between 0800 and 1800h, using standard rat operant chambers (Med Associates, Sandown Scientific, UK) as previously described^22,24^. Operant chambers (30.5×24.1×21.0cm) used for behavioural testing were housed inside a light-resistant and sound-attenuating box. They were equipped with two retractable response levers positioned on each side of the centrally located food magazine. The magazine had a house light (28V, 100mA) located above it. An audio generator (ANL-926, Med Associates, Sandown Scientific, UK) produced tones that were delivered to each chamber via a speaker positioned above the left lever. Operant chambers and audio generators were controlled using K-Limbic software (Conclusive Solutions Ltd., UK).

### Judgement bias training

Animals were trained and tested using a high versus low reward version of the judgement bias task as previously reported^22,24^. Rats were first trained to associate one tone (2kHz at 83dB rats, designated high reward) with a high value reward (four 45mg reward pellets; TestDiet, Sandown Scientific, UK) and the other tone (8kHz at 66dB, designated low reward) with a low value reward (one 45mg reward pellet) if they pressed the associated lever (either left or right, counterbalanced across rats) during the 20s tone (see Figure S1 for a detailed depiction of the task). Unless otherwise specified in Table S1, response levers were extended at the beginning of every session and remained extended for the duration of the session (maximum one hour for all session types). All trials were self-initiated via a head entry into the magazine, followed by an intertrial interval (ITI), and then presentation of the tone. Pressing the incorrect lever during a tone was punished by a 10 s timeout, as was an omission if the rat failed to press any lever during the 20 s tone. Lever presses during the ITI were punished by a 10 s timeout. During a timeout, the house light was illuminated, and responses made on levers were recorded but had no programmed consequences.

Animals underwent a graduated training, and were required to meet criteria for at least two consecutive days before progressing to the next stage. Training stages were as follows:

1. Magazine training: tone played for 20 s followed by release of one pellet into magazine. Criteria: 20 pellets eaten for each tone frequency.
2. Tone training: response on lever during tone rewarded with one pellet. Only one tone frequency, and one lever available per session. Criteria: > 50 trials completed.
3. Discrimination training: response on correct corresponding lever only during tone rewarded with one pellet. Both tones played (pseudorandomly) and both levers available. Criteria: > 70% accuracy for both tones, < 1:1 ratio of correct:premature responses and no significant difference on any behavioural measures analysed over three sessions.
4. Reward magnitude training: As for discrimination training but 2 kHz tone now rewarded with four pellets, 8 kHz tone rewarded with one pellet. Criteria: as for discrimination training but with > 60% accuracy for both tones.

All training sessions consisted of a maximum of 100 trials. Table S1 contains full details of training stages and criteria used. Rats were considered trained when they maintained stable responding for three consecutive days. This was after a maximum of 29 sessions for cohort 1, 25 sessions for cohort 2, and 25 sessions for cohort 3 (see Table S1 for details of session numbers for each training stage).

### Judgement bias testing

Baseline sessions (100 trials: 50 high and 50 low reward tones; presented pseudorandomly, for details see Table S1) were conducted on Monday and Thursday. Probe test sessions (120 trials: 40 high reward, 40 low reward, and 40 ambiguous midpoint tones that were 5kHz at 75dB; pseudorandomly, for details see Table S1) were conducted on Tuesday and Friday. The midpoint tone was randomly reinforced whereby 50% of trials had outcomes as for the high reward tone, and 50% had outcomes as for the low reward tone. This was to ensure a specific outcome could not be learnt, and to maintain responding throughout the experiments (see Figure S1 and Table S1 for a detailed description of how this was implemented). Cohort 1 were used to test the effect of acute systemic treatments with putative RAAD and other NMDA receptor antagonists. Cohort 2 were made up of two groups of eight rats that had previously been used as control animals in another experiment (data not shown) and were then used for the extension of doses of ketamine and PCP. Cohort 3 were used for mPFC infusion experiments. For further details of the different treatments received by each cohort see Table S2.

### Study 1: the effect of acute, systemic treatments with RAADs and NMDA receptors antagonists on judgement bias

#### Experimental design

Each study used a within-subject fully counterbalanced drug treatment schedule (see Table S2 for details of individual treatments). Each animal received all treatments in a counter-balanced design with drug doses separated by a minimum of 72 hrs and at least a one-week drug free period between different treatments. There is the potential for compensatory changes to develop due to repeated testing and the drug treatments, but these are minimised by managing washout periods and also recording and analysing the animals’ baseline data in between drug studies. Results from the baseline sessions are given in Tables S3-S6 and do not indicate any changes in baseline performance due to drug administration for any of the cohorts over time. All drugs were given by intraperitoneal injection using a low-stress, non-restrained technique^35^. Ketamine^¥^ (Sigma-Aldrich, UK), scopolamine^§^ (Tocris, UK), lanicemine^¥^ (Sigma Aldrich, UK), memantine^¥^ (Tocris, UK), MK-801^§^ (Tocris, UK) and PCP^¥^ (Sigma Aldrich, UK) were dissolved in 0.9% sterile saline and given 30^§^ or 60^¥^ minutes prior to testing. CP-101,606 (Experiment 1: Sigma Aldrich, UK; Experiment 2: Boehringer Ingelheim GmbH) was dissolved in 5% DMSO, 10% cremaphor and 85% sterile saline and given 60 minutes prior to testing. Drug doses were selected based on previous rodent behavioural studies^24,36^. Doses for ketamine and PCP were chosen to extend the range of doses tested in this task e.g. higher doses of ketamine and lower doses of PCP were used than previously^24^. For all studies, the experimenter was blind to drug dose. The order of testing for each cohort is displayed in Table S2.

### Study 2: mPFC cannulation and infusions

#### mPFC cannulation

To localize the site and mechanism of action of RAAD drugs, rats were implanted with mPFC guide cannula. Rats were anesthetised with isoflurane/O2 and secured in a stereotaxic frame. Bilateral 32-gauge guide cannulae (Plastics One, UK) were implanted in the mPFC according to the stereotaxic coordinates: anteroposterior +2.7mm, lateral ±0.75mm and dorsoventral −2.0mm from bregma^37^. The cannulae were secured to the skull with gentamicin bone cement (DePuy CMW, UK) and stainless steel screws (Plastics One, UK). Animals received long acting local anaesthetic during surgery, and after surgery the animals were housed individually for 2-3 hours then allowed 10-13 days recovery in normal paired housing conditions. Following the recovery period, rats underwent one week of baseline sessions to re-establish performance. Following this, one week of probe testing was carried out to check that judgement of the ambiguous tone had not altered after surgery. Based on this, another two weeks of probe testing (4 test sessions) was then conducted.

#### Systemic ketamine

Following this, an acute systemic treatment with ketamine was given as a positive control manipulation to ensure that bias could still be manipulated post-surgery. This study was a within-subject fully counterbalanced design, with two treatments (see Table S2, top row of section 3), with the experimenter blind to drug dose. Ketamine (1.0 mg/kg, Sigma Aldrich, UK) was dissolved in 0.9% sterile saline vehicle (0.0 mg/kg) and was given by intraperitoneal injection using a low-stress, non-restrained technique^35^ 60 minutes prior to testing.

#### Infusion Procedure

Rats were then used for mPFC infusion experiments. For details of the infusion procedure. Rats were habituated to the infusion procedure during one session where animals were lightly restrained and the cannula dummy removed and then replaced. In a second habituation session animals were gently restrained while the cannula dummy was removed and a 33-gauge bilateral injector extending 2.5mm beyond the length of the guide cannula was inserted into the mPFC. This was left in place for two minutes, but no infusion occurred. During experimental infusions, the rats were gently restrained while the cannula dummy was removed and the injector inserted. The injector was left in place for 1 min prior to infusions of vehicle or drug (1.0 μl total volume) over 2 minutes. The injector was left in place for a further 2 minutes to allow diffusion of the drug into the tissue surrounding the injector, and then the injector was removed and the dummy replaced. The ambiguous probe test session occurred 5 minutes after the dummy was replaced.

#### Infusion experiments

In the first infusion experiment vehicle (sterile phosphate-buffered saline (PBS); 0.0μg/μl), ketamine (1.0μg/μl), muscimol (0.1μg/μl) or scopolamine (0.1μg/μl), all dissolved in sterile PBS, were infused intracerebrally into mPFC 5 minutes before testing. Following this, CP-101,606 (1.0μg/μl in the first study, 3.0μg/μl in the second study) was dissolved in 10% 2-hydroxypropyl-cyclodextrin and 90% PBS and tested. All experiments used a within-subject fully counterbalanced design for drug treatments, with the experimenter blind to treatment. Drug doses were chosen based on the results from acute, systemic treatments (see Table S2).

#### Histology

Following the completion of mPFC infusions, rats were killed and brains were fixed and processed for histology. Rats were anesthetised with a lethal dose of sodium pentobarbitone (0.5 ml Euthatal, 200 mg/ml, Genus Express, UK) and perfused via the left ventricle with 0.01M PBS followed by 4% paraformaldehyde (PFA). The brains were removed and post-fixed in 4% PFA for 24 hours. Prior to being cut, brains were transferred to 30% sucrose in 0.1M PBS and left for 2 days until brains were no longer floating. Coronal sections were cut at 40 μm on a freezing microtome and stained with Cresyl Violet. Locations of the injector tip positions in the mPFC were mapped onto standardised coronal sections of a rat brain stereotaxic atlas^37^ (Figure 3).

### Data and statistical analysis

Sample size was estimated based on our previous studies using the JBT^22,24^ but with a more conservative effect size as we were looking at acute rather than chronic effects and expected to see greater variation in mPFC infusion studies. Changes in judgement bias should occur without effects on other variables and therefore strict inclusion criteria were established to reduce any potential confound in the data analysis. Only animals which maintained more than 60% accuracy for each reference tone, and less than 50% omissions were used for analysis.

Cognitive bias index (CBI) was used as a measure of judgement bias in response to the midpoint tone. CBI was calculated by subtracting the proportion of responses made on the low reward lever from the proportion of responses made on the high reward lever. This created a score between −1 and 1, where negative values represent a negative bias and positive values a positive bias. Change from baseline in CBI was then calculated for all experimental manipulations as follows: vehicle (0.0mg/kg) probe test CBI - drug dose probe test CBI. This was calculated to take into account individual differences in baseline bias, and to make directional changes caused by drug treatments clearer. To provide a value for vehicle probe test sessions for this measure, the population average for the vehicle (0.0mg/kg) probe test was taken away from each individual rats’ CBI score for the same session. This allowed this measure to be analysed with repeated measures analysis of variance (rmANOVA) with session as the within-subjects factor for drug studies with more than two treatments, or paired samples t-test for studies with only two treatments. The raw data for CBI is included for all drug treatments in Figures S2-S3.

Response latency and accuracy, omissions and premature responses were also analysed (see Table S7 for details of these). These measures were analysed with rmANOVAs with session and tone as the within-subjects factors. Paired t-tests were performed as post-hoc tests if significant effects were established. Huynh-Feldt corrections were used to adjust for violations of the sphericity assumption, and Sidak correction was applied for multiple comparisons. All statistical tests were conducted using SPSS 24.0.0.2 for Windows (IBM SPSS Statistics) with α=0.05. Results are reported with the ANOVA F-value (degrees of freedom, error) and *p*-value as well as any post-hoc *p*-values. All graphs were made using Graphpad Prism 7.04 for Windows (Graphpad Software, USA).

## Results

### Study 1: The effect of acute, systemic treatment with RAADs and selected NMDA receptor antagonists

#### CP-101,606

One animal was excluded in experiments 1 and 2 as accuracy criteria was not met on the vehicle session. In the initial dose response study, CP-101,606 treated animals did not overall show any change in CBI (no main effect of drug dose (*F*_2.237,31.323_=0.811, *p*=0.495). However, compared to baseline performance a one-sample t-test for the highest dose (3.0mg/kg) suggested that there may be small positive shift in CBI (one sample t-test: *p*=0.038; Figure 1A). In the second experiment we tested a higher dose of CP-101,606 (6.0mg/kg), which resulted in a positive bias relative to vehicle treatment (paired samples t-test: *p*=0.027; Figure 1A). In experiment 1, 3.0mg/kg CP-101,606 also caused a decrease in response latency (main effect of session: *F_3,42_*=4.858, *p*=0.005, post-hoc: *p*=0.027; Table 1). There were no effects on other behavioural measures in experiment 1 (Table 1). In experiment 2, CP-101,606 (6.0mg/kg) caused response latencies to decrease (main effect of session: *F_1,14_*=27.396, *p*<0.001; Table 1). This dose had no effect on accuracy for the reference tones (Table 1), but did increase premature responses (paired samples t-test: *p*=0.001), and reduced omissions (main effect of session: *F_1,14_*=10.506, *p*=0.006; Table 1).

**Figure 1.**
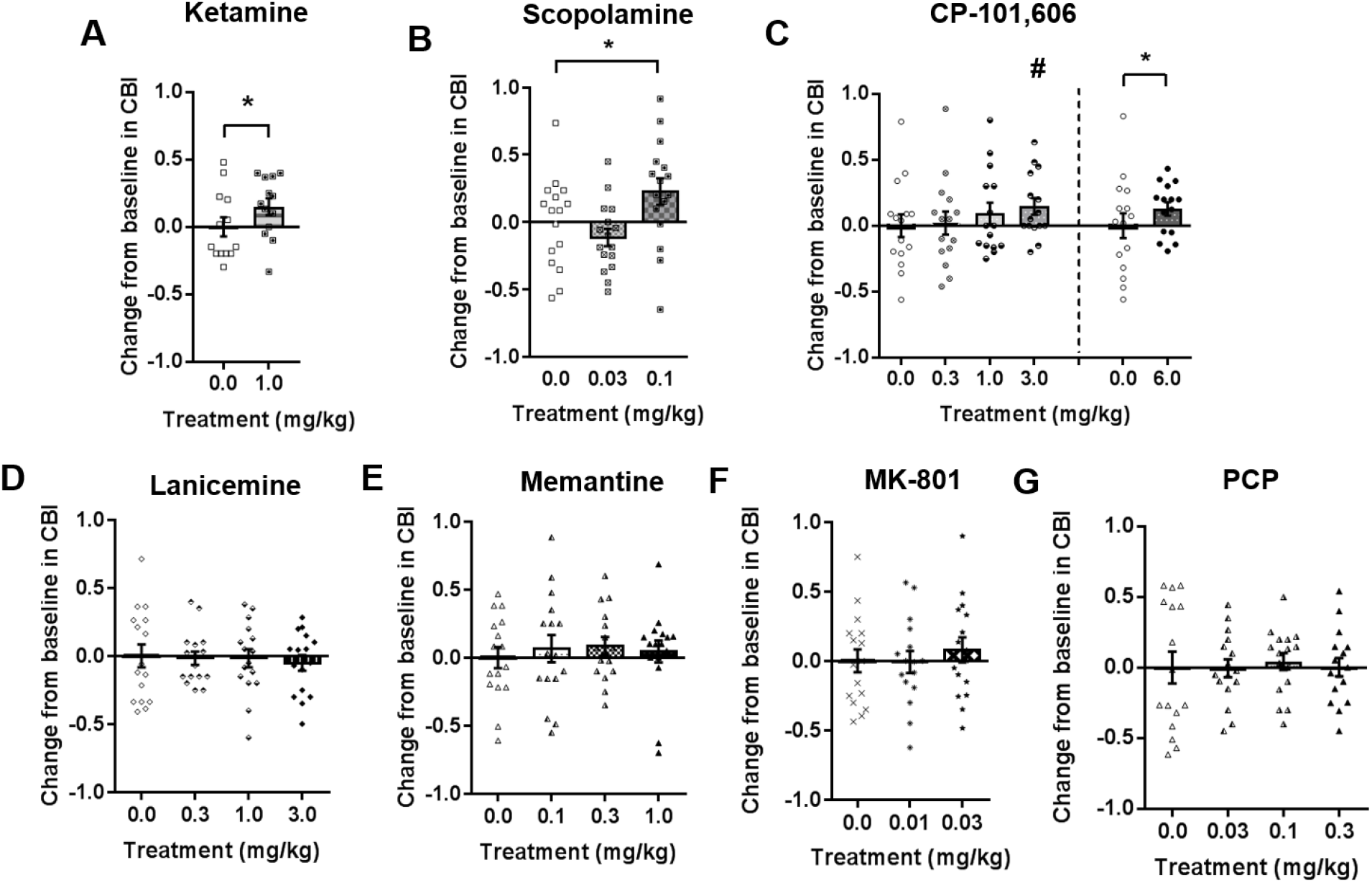
The effect of acute treatment with rapid acting antidepressant drugs and NMDA receptor antagonists on judgement bias of the midpoint ambiguous tone. Ketamine (0.0, 1.0 mg/kg; n = 13), scopolamine (0.0, 0.03, 0.1 mg/kg; n = 16), CP-101,606 (Expt 1: 0.0, 0.3, 1.0, 3.0 mg/kg, n = 15; Expt 2: 0.0, 6.0 mg/kg, n = 15), lanicemine (0.0, 0.3, 1.0, 3.0 mg/kg; n = 16), memantine (0.0, 0.1, 0.3, 1.0 mg/kg; n = 16) and MK-801 (0.0, 0.01, 0.03 mg/kg; n = 16 were administered acutely by intraperitoneal injection prior to testing on the judgement bias task. (A) Replicating previous studies, ketamine (1.0 mg/kg) positively changed CBI. (B) Scopolamine (0.1 mg/kg) also caused a positive change from baseline in CBI. (C) In experiment 1, there was no overall effect of CP-101,606 on change in CBI. A positive change was seen in experiment 2 with a higher 6.0 mg/kg dose. (D-G) Lanicemine, memantine, MK-801 and low doses of PCP did not induce a change in CBI for the midpoint tone at the doses tested. Data shown and represent mean ± SEM (bars and error bars) overlaid with individual data points for each rat. Dashed line (panel C) indicates separate, counterbalanced experiments. \**p* < 0.05; ^#^*p* < 0.05 for a one-sample t-test for 3.0 mg/kg CP-101,6060 only (comparison to a test-value of zero representing a change in CBI for that drug only from baseline). CP-101,606, ketamine, lanicemine, memantine, PCP: 60 min pre-treatment; scopolamine, MK-801: 30 min pre-treatment

**Table 1.** Data for behavioural measures from the JBT.

| Drug / Session | Dose (mg/kg) / Week | Behavioural Measure |  |  |  |  |  |  |  |  |
| --- | --- | --- | --- | --- | --- | --- | --- | --- | --- | --- |
|  |  | Response latency |  |  | % Accuracy |  | % Omissions |  |  | % Premature |
|  |  | HT | MT | LT | HT | LT | HT | MT | LT |  |
| Ketamine | 0.0 | 1.62 ± 0.12 | 3.23 ± 0.22 | 3.80 ± 0.36 | 97.88 ± 0.62 | 83.06 ± 1.96 | 0.00 ± 0.00 | 0.58 ± 0.30 | 1.15 ± 0.36 | 12.44 ± 1.94 |
|  | 1.0 | 1.77 ± 0.13 | 3.38 ± 0.29 | 4.05 ± 0.44 | 97.69 ± 9.57 | 83.54 ± 2.48 | 0.20 ± 0.20 | 0.38 ± 0.26 | 0.78 ± 0.34 | 14.09 ± 3.07 |
| CP-101,606 | 0.0 | 2.19 ± 0.18 | 3.74 ± 0.39 | 4.26 ± 0.32 | 97.82 ± 0.77 | 84.87 ± 1.74 | 0.38 ± 0.26 | 0.88 ± 0.41 | 1.28 ± 0.55 | 8.34 ± 1.24 |
|  | 0.3 | 2.43 ± 0.21 | 3.85 ± 0.36 | 4.54 ± 0.34 | 97.54 ± 0.91 | 85.44 ± 2.39 | 0.00 ± 0.00 | 0.82 ± 0.38 | 1.15 ± 0.47 | 7.42 ± 1.23 |
|  | 1.0 | 2.21 ± 0.15 | 3.48 ± 0.26 | 4.46 ± 0.29 | 98.33 ± 0.72 | 83.49 ± 2.16 | 0.00 ± 0.00 | 0.69 ± 0.41 | 0.83 ± 0.47 | 8.80 ± 1.26 |
|  | 3.0 | *1.97 ± 0.17 | *2.88 ± 0.23 | *3.72 ± 0.27 | 95.83 ± 1.14 | 82.45 ± 2.62 | 0.00 ± 0.00 | 0.17 ± 0.17 | 0.33 ± 0.23 | 12.44 ± 2.58 |
| CP-101,606 | 0.0 | 3.05 ± 0.24 | 4.52 ± 0.27 | 5.38 ± 0.33 | 98.25 ± 0.51 | 86.57 ± 1.70 | 0.70 ± 0.39 | 2.50 ± 0.82 | 2.81 ± 0.80 | 4.24 ± 2.58 |
|  | 6.0 | ***2.32 ± 0.13 | ***3.52 ± 0.17 | ***4.10 ± 0.19 | 97.64 ± 0.87 | 84.37 ± 1.98 | 0.00 ± 0.00 | **0.00 ± 0.00 | **0.72 ± 0.32 | **11.75 ± 1.80 |
| Scopolamine | 0.0 | 2.28 ± 0.17 | 3.84 ± 0.24 | 4.42 ± 0.31 | 97.80 ± 0.64 | 86.56 ± 1.84 | 0.16 ± 0.16 | 0.31 ± 0.21 | 1.09 ± 0.51 | 6.20 ± 0.70 |
|  | 0.03 | ***4.10 ± 0.34 | ***5.52 ± 0.38 | ***6.51 ± 0.40 | **99.50 ± 0.27 | **94.01 ± 1.40 | *7.17 ± 2.17 | ***17.18 ± 3.38 | **18.02 ± 3.60 | 7.31 ± 1.55 |
|  | 0.1 | **4.26 ± 0.38 | **5.54 ± 0.37 | **6.27 ± 0.51 | 95.00 ± 1.76 | 85.26 ± 3.93 | *22.85 ± 3.79 | ***27.98 ± 3.91 | **35.92 ± 6.05 | *14.52 ± 3.85 |
| Lanicemine | 0.0 | 2.42 ± 0.22 | 3.80 ± 0.31 | 4.72 ± 0.45 | 96.60 ± 0.82 | 86.24 ± 1.81 | 0.00 ± 0.00 | 3.16 ± 2.21 | 2.93 ± 1.89 | 7.17 ± 1.11 |
|  | 0.3 | 2.23 ± 0.17 | 3.60 ± 0.24 | 4.40 ± 0.29 | 98.13 ± 0.63 | 85.01 ± 1.89 | 0.00 ± 0.00 | 0.31 ± 0.21 | 1.26 ± 0.95 | 9.48 ± 1.36 |
|  | 1.0 | 2.40 ± 0.22 | 4.20 ± 0.32 | 4.79 ± 0.40 | 96.41 ± 1.58 | 85.45 ± 2.15 | 0.00 ± 0.00 | 3.54 ± 1.99 | 4.03 ± 2.76 | 6.68 ± 1.03 |
|  | 3.0 | 2.55 ± 0.21 | 3.89 ± 0.28 | 4.81 ± 0.37 | 98.58 ± 0.57 | 86.74 ± 1.71 | 0.16 ± 0.16 | 1.56 ± 1.25 | 2.81 ± 1.72 | 8.18 ± 1.67 |
| Memantine | 0.0 | 2.38 ± 0.15 | 4.12 ± 0.21 | 4.94 ± 0.23 | 97.77 ± 0.60 | 83.96 ± 1.52 | 0.19 ± 0.19 | 1.16 ± 0.44 | 2.07 ± 1.03 | 7.98 ± 0.93 |
|  | 0.1 | 2.48 ± 0.18 | 4.15 ± 0.23 | 4.65 ± 0.30 | 97.45 ± 0.82 | 84.75 ± 1.81 | 0.16 ± 0.16 | 2.30 ± 0.85 | 1.09 ± 0.45 | 8.45 ± 1.25 |
|  | 0.3 | 2.42 ± 0.18 | 3.84 ± 0.25 | 4.53 ± 0.30 | 97.49 ± 0.65 | 81.21 ± 1.79 | 0.16 ± 0.16 | 0.63 ± 0.36 | 2.34 ± 0.84 | 9.25 ± 1.40 |
|  | 1.0 | 2.40 ± 0.16 | 3.90 ± 0.23 | 4.50 ± 0.28 | 97.66 ± 0.58 | 84.98 ± 1.64 | 0.00 ± 0.00 | 1.56 ± 0.75 | 2.03 ± 1.25 | 8.38 ± 1.24 |
| MK-801 | 0.0 | 2.51 ± 0.15 | 4.16 ± 0.28 | 4.72 ± 0.30 | 96.39 ± 1.00 | 85.23 ± 1.78 | 0.16 ± 0.16 | 1.12 ± 0.40 | 1.31 ± 0.54 | 6.66 ± 0.77 |
|  | 0.01 | 2.25 ± 0.14 | 3.80 ± 0.21 | 4.39 ± 0.27 | 96.71 ± 0.78 | 83.05 ± 1.64 | 0.16 ± 0.16 | 0.47 ± 0.25 | 1.09 ± 0.51 | 8.39 ± 1.72 |
|  | 0.03 | 2.34 ± 0.13 | *3.44 ± 0.29 | 4.07 ± 0.35 | 97.19 ± 0.88 | 81.19 ± 2.15 | 0.00 ± 0.00 | 1.41 ± 0.56 | 1.88 ± 0.74 | *10.73 ± 1.64 |
| PCP | 0.0 | 2.23 ± 0.17 | 3.88 ± 0.23 | 4.96 ± 0.42 | 98.28 ± 0.50 | 84.89 ± 1.88 | 0.00 ± 0.00 | 0.47 ± 0.25 | 0.47 ± 0.34 | 6.46 ± 0.98 |
|  | 0.03 | 2.22 ± 0.18 | 3.99 ± 0.29 | 5.11 ± 0.42 | 98.59 ± 0.56 | 87.44 ± 1.70 | 0.00 ± 0.00 | 1.25 ± 0.60 | 2.03 ± 1.08 | 5.72 ± 0.75 |
|  | 0.1 | 2.25 ± 0.18 | 3.81 ± 0.29 | 4.60 ± 0.30 | 99.07 ± 0.31 | 86.37 ± 1.87 | 0.31 ± 0.21 | 0.94 ± 0.39 | 1.41 ± 0.94 | 5.63 ± 0.89 |
|  | 0.3 | 2.24 ± 0.20 | 3.93 ± 0.37 | 4.99 ± 0.44 | 98.75 ± 0.51 | 86.78 ± 2.34 | 0.57 ± 0.57 | 1.34 ± 0.73 | 4.11 ± 1.70 | 6.77 ± 1.41 |
| Pre-Surgery |  | 2.05 ± 0.14 | 3.72 ± 0.22 | 4.47 ± 0.33 | 95.80 ± 0.69 | 83.07 ± 2.41 | 0.51 ± 0.30 | 1.50 ± 0.48 | 2.54 ± 0.77 | 14.80 ± 2.30 |
| Post-Surgery | 1 | 1.99 ± 0.15 | 3.82 ± 0.23 | 4.64 ± 0.36 | 95.89 ± 1.00 | 85.62 ± 1.95 | 0.00 ± 0.00 | 0.93 ± 0.31 | 2.18 ± 0.59 | 11.88 ± 1.53 |
|  | 2 | 1.81 ± 0.12 | 3.81 ± 0.31 | 4.29 ± 0.44 | 97.17 ± 0.71 | 84.15 ± 1.45 | 0.17 ± 0.11 | 0.67 ± 0.24 | 2.25 ± 0.62 | 13.33 ± 1.44 |
|  | 3 | 1.77 ± 0.16 | 3.43 ± 0.24 | 4.19 ± 0.40 | 96.58 ± 0.59 | 81.16 ± 2.43 | 0.08 ± 0.08 | 1.00 ± 0.28 | 1.25 ± 0.39 | 11.97 ± 1.70 |

Behavioural data are presented as mean ± SEM. Cells marked in bold indicate a significant difference for that measure, for that tone (compared to vehicle 0.0 mg/kg session for that drug) from a significant dose\*tone interaction. Cells highlighted in grey indicate where a main effect of dose was found, indicating a difference for that drug treatment not specific to tone. \* / light grey shading: $p < 0.05$ , \*\* / medium grey shading: $p < 0.01$ , \*\*\* / dark grey shading: $p < 0.001$ .

#### Scopolamine

The highest dose tested (0.3mg/kg) had to be excluded from the analysis as most rats did not complete sufficient trials. Scopolamine (0.1mg/kg) induced a positive bias (main effect of session: *F_2,30_*=6.739, *p*=0.004, post-hoc: *p*=0.035; Figure 1B). This dose of scopolamine (0.1mg/kg) also increased response latencies (main effect of session: *F_2,30_*=17.263, *p*<0.001, post-hoc: *p*=0.001; Table 1), increased premature responding (main effect of session: *F_1.355,20.330_*=4.387, *p*=0.039, post-hoc: *p*=0.047; Table 1), and increased omissions for all tones (significant session*tone interaction: *F_2.343,35.150_*=4.739, *p*=0.011, main effect of session: *F_2,30_*=24.257, *p*<0.001, post-hoc: *p*s<0.001; Table 1). The lower dose also caused response latencies to increase (post-hoc: *p*<0.001; Table 1), accuracy to increase (main effect of session: *F_1.605,24.069_*=8.558, *p*=0.003, post-hoc: *p*=0.002; Table 1), and omissions to increase for all tones (post-hoc: *p*s≤0.019; Table 1).

#### Ketamine

In the rats who had undergone mPFC cannulation surgery, ketamine (1.0mg/kg) caused a positive change in CBI (paired samples t-test: *p*=0.033; Figure 1C), as has been seen previously^24^. Ketamine did not alter any other behavioural measures (Table 1).

#### Lanicemine

None of the doses of lanicemine tested caused a change in CBI (Figure 1D). This drug also had no effect on any other behavioural measures (Table 1).

#### Memantine

Memantine did not cause any change in CBI at the doses tested (Figure 1E). There was also no effect on other behavioural measures (Table 1).

#### MK-801

MK-801 did not change CBI (Figure 1F). The highest dose of MK-801 tested (0.03mg/kg) decreased response latency for the midpoint tone (main effect of session: *F_2,30_*=3.843, *p*=0.033, and trend towards session*tone interaction: *F_4,60_*=1.332, *p*=0.082, post-hoc: *p*=0.013; Table 1) and also had a tendency to cause premature responses to increase (trend towards main effect of session: *F_1.565,23.480_*=2.915, *p*=0.085, post-hoc: *p*=0.039; Table 1). There was no effect on accuracy for the reference tones or percentage omissions.

#### High-dose ketamine

In experiment 2 (25mg/kg ketamine) one rat was excluded for failure to complete sufficient trials. In experiments 1 and 2, ketamine (10mg/kg and 25mg/kg respectively) did not change CBI (Figure 2A). In both experiments these higher doses did alter all other behavioural measures. There was an increase in response latency across all three tones for both 10mg/kg (session*tone interaction: *F_2,30_*=7.323, *p*=0.003, post-hoc: *p*s<0.001 for all tones; Figure 2B), and 25mg/kg ketamine (session*tone interaction: *F_2,28_*=4.686, *p*=0.018, post-hoc: *p*s≤0.002 for all tones; Figure 2B). Both doses decreased premature responses (paired samples t-tests: 10mg/kg – *p*=0.005, 25mg/kg – *p*=0.006; Figure 2C). Ketamine also improved accuracy for the low reward tone only in experiment 1 (10mg/kg: main effect of session: *F_1,15_*=8.774, *p*=0.010, and trend towards session*tone interaction: *F_1,15_*=3.665, *p*=0.075, post-hoc: *p*=0.022; Figure 2D) and experiment 2 (25mg/kg: trend towards main effect of session: *F_1,14_*=4.500, *p*=0.052, and session*tone interaction: *F_1,14_* = 5.513, *p*=0.034, post-hoc: *p*=0.033; Figure 2D). In both experiments, there was an increase in omissions for all three tones (experiment 1, 10mg/kg: session*tone interaction: *F_1.401,21.021_*=5.662, *p*=0.018, post-hoc: high reward tone – *p*=0.015, midpoint tone: *p*=0.003, low reward tone: *p*=0.010; experiment 2, 25mg/kg: session*tone interaction: *F_1.368,19.150_*=11.964, *p*=0.001, post-hoc: high reward tone – *p*=0.003, midpoint tone – *p*<0.001, low reward tone – *p*=0.001; Figure 2E).

**Figure 2.**
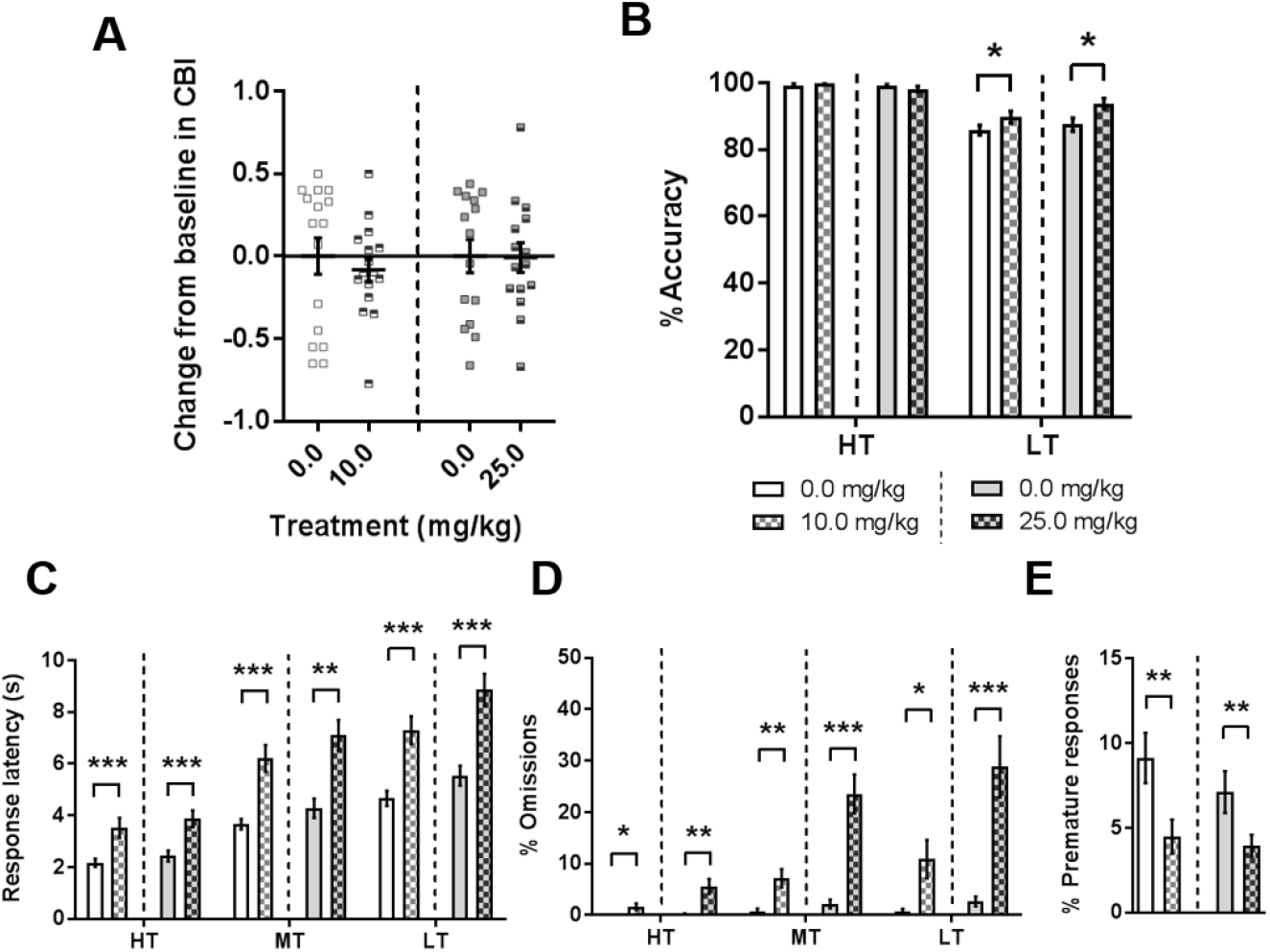
Behavioural data from the judgement bias task following acute treatment with high doses of ketamine. Acute doses of ketamine (Expt 1: 0.0, 10.0 mg/kg, n = 16; Expt 2: 0.0, 25.0 mg/kg, n = 16) were administered by intraperitoneal injection to measure their effect on judgement bias. (A) Neither high dose of ketamine caused a change in interpretation of the midpoint tone. (B) Both doses of ketamine increased accuracy for the low tone. (C) Both doses of ketamine increased response latencies across all three tones. (D) Omissions were increased across all three tones following both ketamine doses. (E) High doses of ketamine (10.0, 25.0 mg/kg) decreased premature responding. \*\*\**p* < 0.001, \*\**p* < 0.01, \**p* < 0.05. Data represent mean ± SEM (panels B-E) with individual data points overlaid for each rat (panel A). Dashed lines indicate separate, counterbalanced experiments. 60 min pre-treatment. HT - high reward tone; MT - midpoint tone; LT - low reward tone.

#### Low dose PCP

Doses of PCP (0.03, 0.1, 0.3mg/kg) that were lower than those previously tested^24^ did not cause any change in CBI (Figure 1G). There was also no effect on any other behavioural measures (Table 1).

#### Analysis of performance split over session

In addition to the analyses above we also compared performance for the first and last 20 probe trials in order to check whether animals’ performance changed within a session during these randomly reinforced trials. Analysis of the data for doses of ketamine (1.0mg/kg), CP101606 (6.0mg/kg) and scopolamine (0.1mg/kg) which change CBI did not find any evidence of differences across the session between vehicle or drug treatments based on this analysis (see Figure S4).

### Study 2: mPFC infusions of drugs shown to cause positive judgement biases

Two rats were excluded in cohort 3: one rat did not meet accuracy criteria for any probe (or baseline) session following the second drug infusion; and after the end of testing another animal was found to have an incorrect cannula placement. Therefore, both were excluded retrospectively from the entire study. Compared to pre-surgery performance, the CBI of rats became more negative after surgery, and this was stable across testing over three weeks (main effect of week: *F_3,42_*=6.335, *p*=0.001, post-hoc: *p*s≤0.011; Figure 3A). There were no differences in response latencies, premature responses, accuracies for reference tones or omissions before compared to after surgery (Table 1). The change in CBI occurred before infusions and seemed to be a response to the surgical intervention potentially causing a more negative affective state. We found no evidence of tissue damage in the area surrounding the cannula post-mortem, so it is unlikely that this was a result of trauma. We think it is not surprising that undergoing surgery and having to adapt to intracerebral cannula could cause a permanent negative change in affect. It is exactly this sort of affective state change that judgement bias assays have been developed to detect (for example see Bethell^38^, and Baciadonna & McElligott^39^ for reviews summarising how judgement bias tasks can be used as measure of animal welfare).

**Figure 3.**
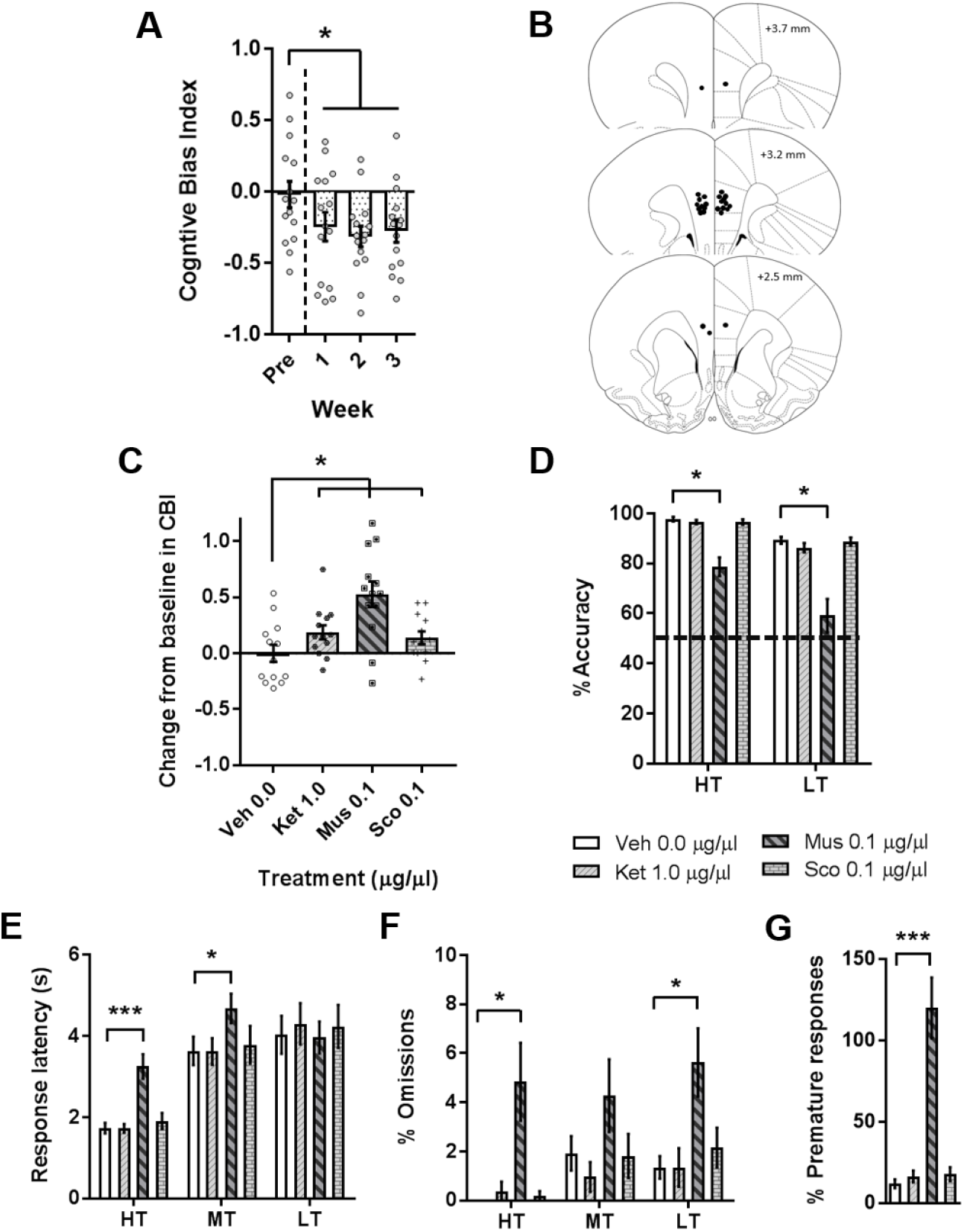
D*ara from mPFC cannulated rats on the judgement bias task*. Probe tests with no experimental manipulation were conducted before and after mPFC cannulation surgery to ensure that the surgery itself did not effect performance in the judgement bias task. (A) Cognitive bias index became more negative in the probe tests conducted after surgery. (B) The location of the injector placement was confirmed post-mortem and black dots represent the location of the cannula tip as assessed from Cresyl violet-stained brain sections. Coronal sections are +3.7 mm to +2.5mm relative to bregma (Paxinos and Watson, 1998). (C-G) In the first infusion experiment, ketamine (Ket; 1.0 μg/μl) muscimol (Mus; 0.1 μg/μl), scopolamine (Sco; 0.1 μg/μl) or vehicle (Veh; 0.0 μg/μl; n = 13), were administered by intracerebral infusion into the mPFC to measure the effect on judgement bias. (C) Ketamine, muscimol and scopolamine all caused a positive change in cognitive bias index (CBI) for the midpoint tone. (D) Muscimol decreased accuracy for both reference tones. (E) Muscimol increased response latencies for the high and midpoint tones. (F) For the high and low tones, muscimol increased omissions. (G) Muscimol also increased premature responding. Data represent mean ± SEM (panels A, C-G) with individual data points overlaid for each rat (panel A,C). Black dashed line (panel f) represents 50% accuracy depicting performance at chance. 5 min pre-treatment. \*\*\**p* < 0.001, \**p* < 0.05. HT - high reward tone; MT - midpoint tone; LT - low reward tone.

In the first infusion experiment, ketamine (1.0μg/μl), muscimol (0.1μg/μl) and scopolamine (0.1μg/μl) all induced positive biases (main effect of session: *F_3,36_*=7.241, *p*=0.001; post-hoc: ketamine – *p*=0.012, muscimol – *p*=0.001, scopolamine – *p*=0.032 Figure 3C). The effect of PFC infusion of ketamine or scopolamine was specific to CBI, as these drugs had no effect on other behavioural measures (Figure 3D-G), unlike muscimol infusions which caused changes to all other behavioural measures. There was an increase in response latency (session*tone interaction: *F_6,72_*=4.181, *p*=0.001) for the high reward (post-hoc: *p*<0.001) and midpoint tone (*p*=0.028; Figure 3D), and a large increase in premature responses to over 100% (main effect of session: *F_1.151,13.809_*=33.784, *p*<0.001, post-hoc: *p*<0.001; Figure 3E). Muscimol also caused accuracy to decrease (main effect of session: *F_1.181,14.172_*=43.775, *p*<0.001, post-hoc: *p*≤0.001; Figure 3F). For the low reward tone, this reduction was so great that rats were no longer performing any better than chance (one-sample t-test against a test value of 50%: *p*=0.197; Figure 3F). Omissions increased following muscimol infusion (main effect of session: *F_1.338,16.057_*=10.418, *p*=0.003, post-hoc: *p*=0.007; Figure 3G).

In the experiments testing the effect of CP-101,606 mPFC infusion, in experiment 1 the lower dose (1.0μg/μl) did not alter CBI (Figure 4A), but in experiment 2, the higher dose (3.0μg/μl) induced a positive bias (paired samples t-test: *p*=0.043; Figure 4A). In experiment 1, CP-101,606 (1.0μg/μl) caused an increase in response latency (main effect of session: *F_1,12_*=5.064, *p*=0.044; Figure 4B) but had no other behavioural effects (Figure 4C-E). In experiment 2, 3.0μg/μl CP-101,606 did not have any effects on other behavioural measures (Figure 4B-E).

**Figure 4.**
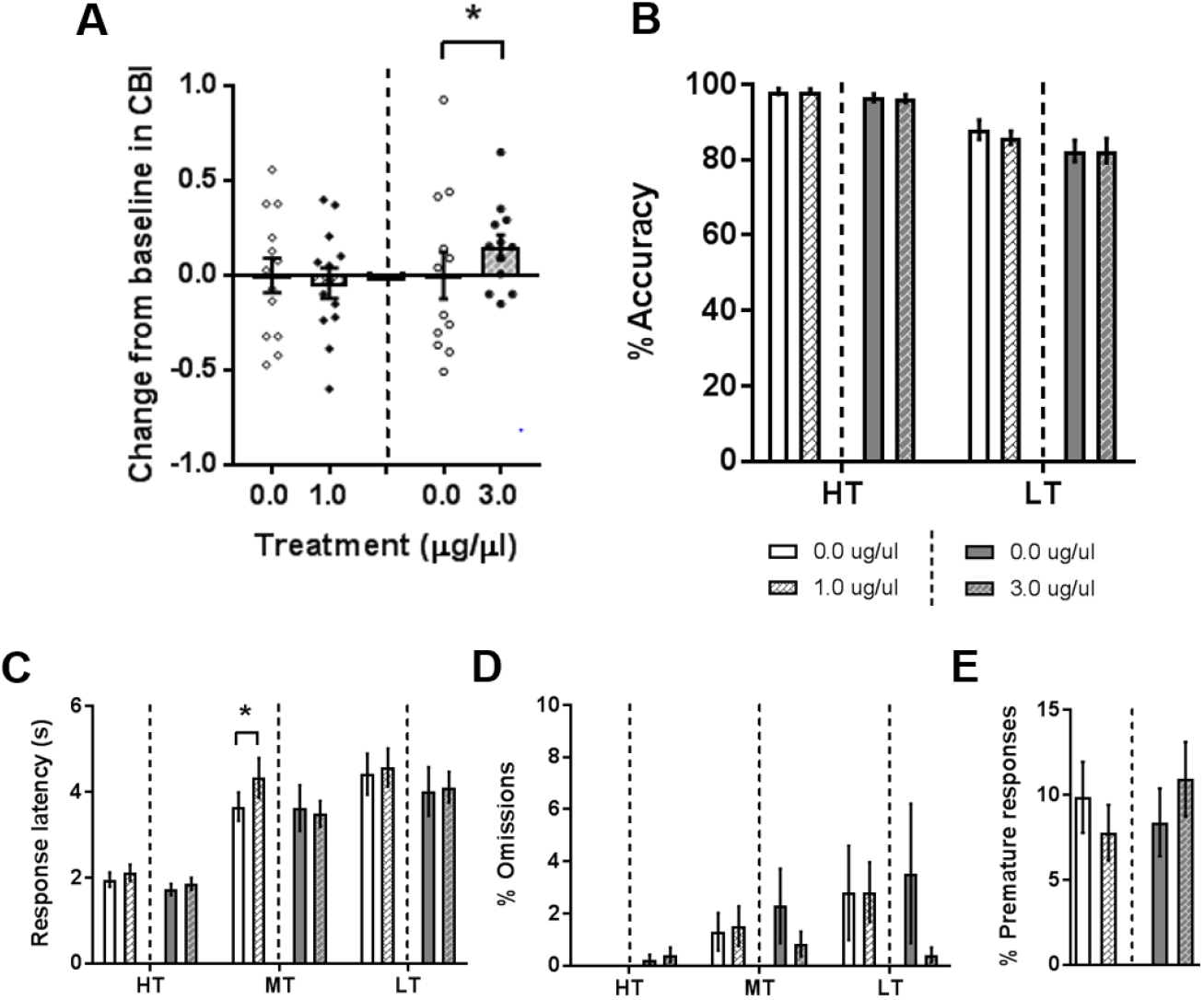
Behavioural data from the judgement bias task following mPFC infusions of CP-101,606. CP-101,606 (Expt 1: 00.0, 1.0 μg/μl, n = 13; Expt 2: 0.0, 3.0 μg/μl, n = 12) was administered by intracerebral infusion in the mPFC to measure the effect on judgement bias. (A) The higher dose of CP-101,606 (3.0 μg/μl) caused a positive change from baseline in CBI. (B) Accuracy was not altered by either dose of CP-101,606. (C) In experiment 1, CP-101,606 (1.0 μg/μl) increased response latency for the midpoint tone. (D/E) There was no effect of either dose on omissions or premature responding, \**p* < 0.05. Data represent mean ± SEM (panels B-E) with individual data points overlaid for each rat (panel A). Dashed lines indicate separate, counterbalanced experiments. 5 min pre-treatment. HT - high reward tone; MT - midpoint tone; LT - low reward tone.

## Discussion

As previously shown^24^, low dose ketamine (1.0mg/kg) had a specific effect on decision-making biases, inducing a positive change in CBI following acute administration. This effect of ketamine was dose dependent, with higher doses having general effects on task performance without changing CBI. The effects of ketamine were recapitulated to some extent by the GluN2B antagonist, CP-101,606 and muscarinic antagonist, scopolamine, but both also had more general effects on other behavioural measures following systemic administration. All three treatments have previously been reported to have RADD effects in clinical trials^3,29,30^, whilst the other NMDA antagonists tested here did not^5,26–28^, and these also failed to induce a change in bias. The mPFC infusions suggest that this brain region is central to the effects of ketamine, scopolamine and CP-101,606. Interestingly, mPFC infusions more specifically altered bias, suggesting other brain regions may contribute to the systemic effects on other behavioural measures. The importance of the mPFC in modulating RAAD effects in neuropsychological tasks is consistent with previous findings in our learning and memory bias assay, the affective bias test^34^. Inactivation of the mPFC with muscimol did positively change bias but animals also exhibited large changes in other behavioural measures. This suggests that the RAADs can modulate activity in this brain region in a more specific way than muscimol, which results in a relatively specific effect on biases in decision-making.

For lanicemine and memantine, the lack of any behavioural effects means there is a possibility that the doses tested were too low. For both treatments the range of doses tested covers the doses that are equivalent to those used humans in clinical trials (lanicemine: 50, 100mg^26^, equivalent to approximately 0.75, 1.5mg/kg; memantine: 5-20mg^5^, equivalent to approximately 0.07-0.3mg/kg), paralleling our effective dose of ketamine (1.0mg/kg, similar to the 0.5mg/kg dose used by Zarate et al.^3^). Although higher doses may yield behavioural effects, these are likely to be due to much higher levels of receptor occupancy than those relevant to the antidepressant effects and may also arise from non-specific actions at other receptors. When testing lower doses of PCP (another NMDA receptor antagonist not known to show RAAD) than previously used^24^, we also failed to see any change in CBI. Conversely, when we tested higher doses of ketamine than those we had previously^24^, doses that are often used to demonstrate antidepressant effects in other preclinical models used to study depression such as the forced swim test (FST)^40^, we failed to see any change in bias, instead only seeing non-specific changes in other behavioural measures. The behavioural profile seen with these higher doses of ketamine (increased response latency and omissions and decreased premature responding) suggests that these doses may be causing locomotor depression or reducing motivation to respond. Higher doses of ketamine have not been found to have antidepressant effects in clinical trials and these data also suggest that rodent studies using these higher doses may not be looking at specific effects. It may be that the lower 1.0mg/kg dose of ketamine can specifically alter decision making biases because they target a specific population and hence modulate a specific circuit. Some studies have suggested that ketamine may act via disinhibition of GABAergic interneurons leading to a glutamate burst which then activates prefrontal glutamate neurons^41^. Overall, the results from systemic administration of different NMDA receptor antagonists lends support to our interpretation that this reward-based judgement bias task can specifically dissociate between drugs that do show RAAD, and those that do not, despite them having similar pharmacology.

The difference in specificity on behavioural effects, whereby ketamine (1.0mg/kg) only positively changes decision-making bias, but both CP-101,606 and scopolamine have other non-specific effects, suggests that 1.0mg/kg ketamine is able to relatively selectively modulate affective bias. The changes in response latencies, omissions and premature responses caused by CP-101,606 and scopolamine suggest that these drugs may also be having effects on other cognitive processes, such as motivation. However, the direction of changes for these drugs are in opposite directions (decreases in response latency and omissions for CP-101,606 but increases in these for scopolamine) despite them both causing positive changes in CBI. This, combined with the lack of change in accuracy for the reference tones, suggest that these non-specific effects cannot fully explain the change in decision-making bias.

The neurobiology underlying the relative specificity of ketamine, CP-101,606 and scopolamine in being able to immediately alter decision-making bias, in contrast to the other NMDA receptor antagonists tested that have not shown these effects, are likely to be due to differences in their mechanisms of action. Our findings add weight to the strong body of evidence suggesting that NMDA receptor antagonism is important for short-term, RAAD effects of these drugs^42^, but suggests that specific modulation of either a specific subtype of the receptor or a sub-population of neurons may be involved. CP-101,606 is selective for the GluN2B NMDA receptor subunit, whilst it has been shown that scopolamine, and more recently ketamine, cause a glutamate burst via blockade of NMDA receptors specifically on GABA interneurons that leads to increased mechanistic target of rapamycin complex 1 signalling, brain-derived neurotrophic factor release and synaptic changes in the PFC^41,43–45^. Further studies would be required to test whether these mechanisms also drive these drugs effects on affective bias.

The infusion studies localise the site of action of this rapid change in decision-making bias to the mPFC, corresponding with brain imaging studies in humans that have also shown ketamine-dependent changes in prefrontal glutamatergic neurotransmission^32,33^. This also matches with previous rodent studies, where using the affective bias test, it has been shown that whilst ketamine does not induce positive biases in learning, it can remediate previously acquired negative biases, an effect which also localises to the mPFC^34^. For CP-101,606 and scopolamine, unlike when given systemically, intracerebral mPFC infusion did not cause any non-specific behavioural changes on the task. This could suggest that these non-specific effects are driven by off-target effects of drug binding in other brain areas, or in the case of scopolamine, the periphery. The localisation of the positive modulation of decision making caused by these drugs to the mPFC provides further support for the hypothesis that this might be mediated through burst firing in the prefrontal cortex, an effect that has recently been shown to cause the activation of downstream pathways thought to be important in the RAAD effects of both ketamine and scopolamine^41,43–45^.

Interestingly, both GABA_A_ receptor agonism (musciol infusion), and NMDA receptor antagonism (ketamine infusion) in the mPFC caused the same qualitative, but not quantitative behavioural change in judgement bias (a positive shift but of different magnitudes), mirroring findings seen previously with intra-infralimbic infusions of muscimol and (R)-CPP on the five choice serial reaction time task, where both drugs increased impulsive responding but by different amounts^46^. It has been suggested that the functional effects of NMDA receptor antagonism may be due to excess extracellular glutamate^47,48^. However, the pronounced, non-specific behavioural effects on other measures seen following muscimol infusion suggests that mechanism of action of the other infusion drugs is more refined than global inhibition of neurotransmission in the mPFC. Previous work in humans and rodents has shown that subcortical and limbic brain regions, such as the amygdala, are important in the neurocircuitry of MDD / depression-related behaviour^49–51^, and a recent study suggests that ketamine may play a critical role in restoring dysfunctional connectivity in these circuits^33^. Furthermore, in rodents, a recent study found that optogenetic activation of pyramidal mPFC neurons containing dopamine receptor D1 caused RAAD-like responses in the forced swim test, and that blockade of these receptors prevented the RADD effects of ketamine^52^. In order to further our understanding of this mechanism, it will be important to investigate the effects of these drugs on different neuronal subtypes within the mPFC, as well as investigating the wider circuitry that is altered by these drugs.

### Final conclusions

This study adds to the evidence that the neuropsychological effects of ketamine are potentially important in its RAAD in patients with MDD, and that these effects in altering affective biases, both in decision-making as demonstrated here, as well as in learning and memory occur at time points (one hour) before major plastic changes arise. It will be important to investigate the neurobiological effects of not just the immediate, RAAD of ketamine, but also the sustained effects by examining how affective biases are altered at longer time points. Furthermore, investigation of the wider circuits involved in this RAAD efficacy will be crucial in revealing the mechanism underlying these actions, which will be important for the development of novel therapeutics. Ketamine (at 1.0mg/kg) seems to have very specific effects on affective bias, which we can capitalise on to better understand the circuits that contribute to these modulations of affective biases that are potentially very important in the cause, perpetuation and treatment of MDD. More detailed circuit analyses are needed including undertaking studies in other brain regions to determine whether ketamine’s effects are specific to the mPFC.

## Funding and Disclosure

This research was funded by an Industrial Partnership Award awarded by BBSRC in collaboration with Boehringer Ingelheim (Grant no: BB/N015762/1) and carried out with intellectual support from Boehringer Ingelheim. ESJR has current or previously obtained research grant funding through PhD studentships, collaborative grants and contract research from Boehringer Ingelheim, Compass Pathways, Eli Lilly, MSD, Pfizer and SmallPharma. The authors declare no conflict of interest.

## Author contributions

CAH performed the research, analysed data, wrote and edited the paper. JMB performed research and analysed data. RA and BH designed the research and edited the paper. ESJR designed the research and wrote and edited the paper.

## Notes

### Competing Interest Statement

The authors have declared no competing interest.

### Summary of Updates

Background has been added to the introduction about judgement bias tasks; further details have been provided in the methods about the task structure, trial format and training, and parts of the discussion have been expanded.

