## Supplementary Figures and Tables for "Role of the medial prefrontal cortex in the effects of rapid acting antidepressants on decision-making biases in rodents"

**Running title: Rapid antidepressant effects on decision-making biases**


**Author names:** Hales CA[1] (PhD), Bartlett JM[1] (BSc), Arban R[2] (PhD), Hengerer B[2] (PhD), Robinson ESJ[1] (PhD)

**Author Affiliations:** [1]School of Physiology, Pharmacology and Neuroscience, Faculty of Biomedical Sciences, University of Bristol, Bristol, BS8 1TD, UK

[2]CNS Diseases Research, Boehringer Ingelheim GmbH & Co. KG, Biberach an der Riss, Germany

**Corresponding author:** Name: Prof. Emma Robinson

Address: School of Physiology, Pharmacology and Neuroscience, Faculty of Biomedical Sciences, Tankards Close, University of Bristol, Bristol, BS8 1TD, UK.

Telephone: (+44)117 3311449

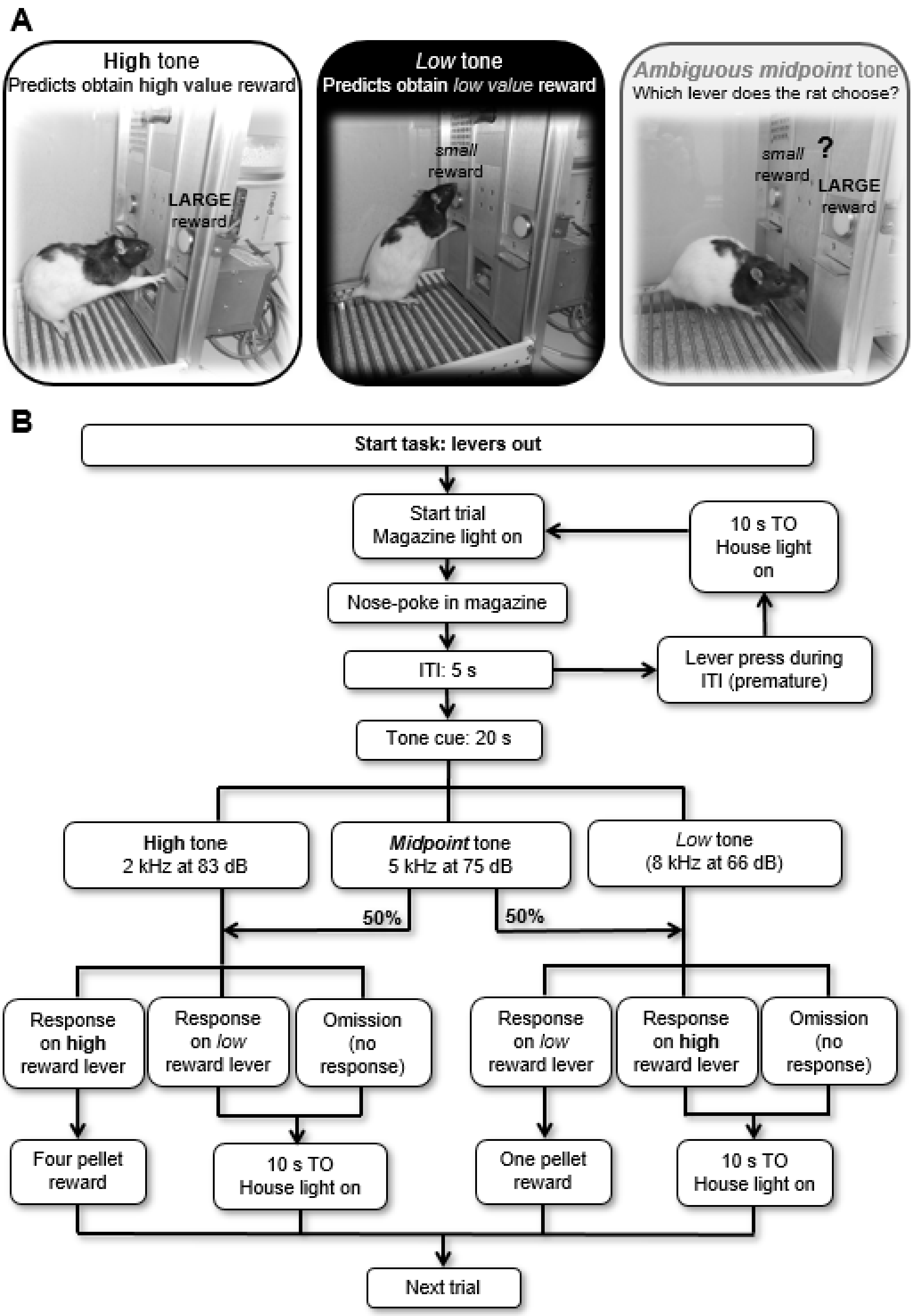


### Figure S1 – Schematic of the JBT and trial structure.

In the judgement bias task (JBT), rats are trained to associate one tone frequency (2 kHz) with a high value reward: i.e. if the rat presses the correct lever (shown as the left lever in (A), but counterbalanced across rats in a cohort) they receive a high value reward (four reward pellets). They also learn to associate a second tone frequency (8 kHz) with receiving a low value reward (one reward pellet; shown in (A) as pressing the right lever during the tone). Judgement bias, or decision making about an ambiguous cue, which is known to be influecned by affective state, can be probed by presenting an ambiguous tone that has a mipdoint frequency between the two reference cues (5 kHz), and recording which lever the rat presses. If the rat is expecting the more positive outcome (indicative of an optimistic judgement bias), then they will more often choose the large reward lever, but if the rat is in a more negative affective state, they will expect the less positive outcome and more often choose the low reward lever, a pessimistic judgement bias. During the task, tones are presented within discrete trials, the format of which is depicted as a flow chart in (B). The task is self-initiated, and so each trial begins only once the rat makes a nosepoke entry into the magazine port. This is followed by a 5 second inter-trial interval (ITI), during which time the rat has to wait and refrain from making a lever press response. If the rat does press a lever, they are punished with a 10 second timeout (TO). The tone cue is presented for a maximum of 20 seconds following the ITI, or until the rat makes a lever press response. The outcome following each lever press depends on which tone was played, and which lever was pressed. Correct lever presses to either reference tone (high or low tones) results in the corresponding reward being delivered to the magazine, whilst incorrect lever presses results in a 10 second TO. This TO also occurs if the rat fails to make any lever press during the 20 second tone presentation (an omission). During TOs, lever presses and magazine entries are recorded but have no consequences, meaning the rat has to wait to be able to beign the next trial. When the midpoint tone is presented, 50% of the time this tone is “classified” by the software as having the same response properties as the high reward tone. I.e., if the rat makes a high reward lever press during a midpoint tone presentation classified in this way, then they will receive a four pellet reward, but will experience the 10 second TO if they make a low reward lever press. Similarly, if the midpoint tone is “classified” as having the same response properties as the low reward tone, then a high reward lever press would result in a TO, whilst a low reward lever press would result in delivery of the small reward. In this way, each lever is only every associated with the same reward outcome (i.e. four pellets for the high reward lever), but the midpoint tone becomes randomly reinforced, and so rats will maintain responding for this tone across multiple trials within a session, whilst being unable to learn a specific reward contigency to associate with the midpoint tone.

**Table S1** - Training stages and required performance criteria for the judgement bias task.

| Stage | Description | Criteria | Sessions required to meet criteria | | |
|---|---|---|---|---|---|
| | | | Cohort 1 | Cohort 2 | Cohort 3 |
| **1 – Magazine training** | Tone (2 kHz only for half the session followed by 8 kHz only for the rest of the session, order counterbalanced across rats) played for 20 s followed by release of one pellet into magazine; 10 s ITI. No levers available. | 20 pellets eaten for each tone frequency | 1 | 1 | 1 |
| **2 – Tone training** | Response on lever during tone (2 kHz or 8kHz only, order counterbalanced across rats) rewarded with one pellet. Lever corresponding to that tone frequency available only. | > 50 trials completed for two consecutive sessions on each tone frequency | 4 | 4 | 4 |
| **3 – Discrim-ination training** | Response on correct corresponding lever only during tone (either 2 kHz or 8 kHz presented pseudorandomly) rewarded with one pellet. Both levers available. Incorrect or omitted trials were repeated (i.e. same tone frequency played) until a correct response occurred. | > 70% accuracy for both tones, no significant differences on analysed behavioural measures over three sessions and < 1:1 ratio of correct:premature responses | 15 | 10 | 10 |
| **4 – Reward magnitude training** | As Stage 3 but response on correct corresponding lever only rewarded with four pellets for high reward tone and one pellet for low reward tone. Both levers available. | As for Stage 3 but with > 60% accuracy for both tones (to allow for biases in responding to reference tones caused by the difference in associated reward magnitude). | 9 | 8-10 | 10 |
| **5 – Baseline session** | Same format as reward magnitude training sessions. | Animals had to show equivalent baseline session performance to pre-drug study baseline sessions (measured by no significant differences pre- and post- on behavioural measures. | - | - | - |
| **6 – Probe sessions** | For reference tones (2 of 8 kHz) response on correct corresponding lever during the tone rewarded with either 4 pellets (2 kHz) or 1 pellet (8 kHz) reward. For ambiguous midpoint tone (5 kHz), random reinforcement was used whereby outcomes for 50% of the trials followed 2 kHz tone trials, whilst the other 50% followed 8 kHz tone trials (see Supplementary Figure XX for further details). There were no repeated trials following incorrect or omissions. | < 60% accuracy for both reference tones, < 50% omissions | - | - | - |

For all training stages, trial structure was as depicted in Figure S1 (except for magazine training which excludes any form of lever press response). Training stages 1-4 were conducted once per day, Monday to Friday, and consisted of a maximum of 100 trials, or lasted for 60 minutes. Where both tones were played (stages 3-4) tone type was equally spilt across the session (50 trials per tone). Baseline sessions also consisted of 100 trials, and were conducted Monday to Friday during baseline weeks between drug studies, and on Monday and Thursday during drug studies. Probe sessions consisted of 120 trials: 40 of each reference tone (2 and 8 kHz) and 40 midpoint tones (5 kHz). Pseudorandom tone presentation was achieved by splitting each training/baseline (probe) session into blocks of 10 (12) trials, within which there were 5 (4) presentations of each tone frequency. Within a block, there could only be a maximum of n-1 consecutive tone presentations (i.e. in baseline sessions, a maximum of 4 consecutive trials of either 2 or 8 kHz). Omitted trials (no lever press during tone presentation; possible in stages 2-4) were punished with a 10 second timeout where the house light was turned on, and the animal was unable to initiate another trial. Incorrect trials (wrong lever for the tone presented; possible in stages 3-4) were also punished with a 10 second timeout with the house light on. Premature trials (lever press during the ITI; possible during stage 2-4) were similarly punished with a 10 second timeout with the house light on. Each new trial had to be self-initiated by the animal by making an entry into the magazine (this was signalled by the magazine light being turned on, and was switched off once animals made the magazine nose poke). Baseline session were the same format as reward magnitude training sessions, with animals required to repeat incorrect or omitted trials. Midpoint tones during the probe session were reinforced as follows: to program random reinforcement within the constraints of the software, the ambiguous midpoint tone was made up of two copies of the 5 kHz tone (75 dB), each of which was programmed to be classed as “correct” for one of the two lever press responses. This meant that the outcome associated with each of the two ambiguous tones could be programmed to be the same as one of the reference tones, hence resulting in random reinforcement. I.e., 50% of the time lever presses for the midpoint tone had outcomes that were the same as the high reward tone (4 pellets or a timeout), whilst 50% of the time lever presses had outcomes that were the same as the low reward tone (timeout or 1 pellet). Responses to either of the “two” midpoint tones were analysed together.

**Table S2** - Summary of treatments used in the different cohorts.

| Cohort | # rats | Acute drug treatment | Doses (mg/kg) |
|---|---|---|---|
| **1** | 16 | Memantine<br>MK-801<br>Lanicemine<br>CP-101,606 (Experiment 1)<br>CP-101,606 (Experiment 2)<br>Scopolamine | 0.0, 0.1, 0.3, 1.0<br>0.0, 0.01, 0.03<br>0.0, 0.3, 1.0, 3.0<br>0.0, 0.3, 1.0, 3.0<br>0.0, 6.0<br>0.0, 0.03, 0.1, 0.3 |
| **2** | 16 | Low dose PCP<br>High dose ketamine (Experiment 1)<br>High dose ketamine (Experiment 2) | 0.0, 0.03, 0.1, 0.3<br>0.0, 10.0<br>0.0, 25.0 |
| **3** | 15[$] | Ketamine (systemic)<br>mPFC infusions (Experiment 1): ketamine, muscimol, scopolamine<br>mPFC infusion: CP-101,606 (Experiment 1)<br>mPFC infusion: CP-101,606 (Experiment 2) | 0.0, 1.0<br>0.0, 1.0, 0.1, 0.1 µg/µl<br>0.0, 1.0 µg/µl<br>0.0, 3.0 µg/µl |

[$]Initial total n number for this manipulation is 15 as one rat had to be euthanised after the first infusion habituation session as dummy cannula could not be removed from the guide.

**Table S3 –** Data for response latency for baseline weeks between drug studies from the JBT.

| Response Latency | | | | | | | | | | | |
|---|---|---|---|---|---|---|---|---|---|---|---|
| | | Day 1 | | Day 2 | | Day 3 | | Day 4 | | Day 5 | |
| Drug | Cohort | HT | LT | HT | LT | HT | LT | HT | LT | HT | LT |
| Memantine | 1 | 2.26±0.17 | 3.68±0.26 | 2.54±0.15 | 4.65±0.28 | 2.71±0.19 | 5.08±0.30 | 2.63±0.17 | 4.59±0.27 | 2.84±0.21 | 4.39±0.33 |
| MK-801 | | 2.52±0.19 | 4.57±0.32 | 2.92±0.15 | 5.61±0.28 | 2.97±0.15 | 5.35±0.23 | 2.82±0.16 | 5.20±0.26 | 3.11±0.22 | 5.72±0.29 |
| Lanicemine | | 2.60±0.15 | 4.81±0.24 | 2.85±0.14 | 5.04±0.25 | 2.81±0.15 | 4.82±0.21 | 2.95±0.18 | 5.26±0.20 | 3.05±0.34 | 5.80±0.43 |
| CP-101,606: low | | 2.58±0.21 | 4.59±0.26 | 2.72±0.15 | 4.94±0.29 | 2.72±0.20 | 4.91±0.26 | 2.48±0.19 | 4.44±0.17 | 2.94±0.21 | 5.42±0.20 |
| CP-101,606: high | | 2.99±0.23 | 5.11±0.31 | 3.15±0.25 | 5.30±0.29 | 3.28±0.24 | 5.84±0.26 | 3.02±0.21 | 5.34±0.30 | 3.68±0.22 | 5.80±0.29 |
| Scopolamine | | 2.62±0.22 | 4.92±0.33 | 2.97±0.19 | 5.23±0.30 | 3.93±0.24 | 5.69±0.32 | 3.03±0.22 | 5.62±0.28 | 4.32±0.71 | 6.16±0.30 |
| PCP | 2 | 2.29±0.13 | 4.52±0.37 | 3.07±0.29 | 6.12±0.43 | 2.88±0.31 | 5.74±0.46 | 2.82±0.29 | 5.59±0.47 | 2.84±0.24 | 5.36±0.46 |
| Ketamine (10) | | 3.54±0.29 | 5.99±0.44 | 4.24±0.40 | 7.17±0.49 | 3.71±0.20 | 6.01±0.32 | 3.73±0.22 | 6.29±0.41 | 3.53±0.29 | 6.12±0.31 |
| Ketamine (25) | | 2.63±0.26 | 4.17±0.47 | 3.21±0.22 | 5.61±0.35 | 3.48±0.24 | 6.42±0.49 | 3.54±0.20 | 6.41±0.31 | 3.62±0.26 | 5.93±0.28 |
| Post-Surgery | 3 | 1.86±0.13 | 3.20±0.26 | 1.97±0.10 | 4.06±0.27 | 2.08±0.11 | 4.33±0.27 | 2.00±0.15 | 4.37±0.24 | 2.49±0.15 | 4.61±0.34 |
| Ketamine (1) | | 2.01±0.15 | 4.03±0.30 | 2.29±0.22 | 4.47±0.42 | 2.34±0.22 | 4.32±0.39 | 2.62±0.21 | 4.60±0.32 | 2.41±0.15 | 4.19±0.31 |
| Infusions 1 | | 2.14±0.24 | 4.33±0.38 | 2.43±0.27 | 4.89±0.37 | 3.02±0.30 | 5.69±0.41 | 2.54±0.25 | 5.19±0.36 | 2.32±0.20 | 4.99±0.34 |
| CP-101,606 infusion: low | | 2.61±0.21 | 5.16±0.35 | 2.70±0.19 | 5.54±0.42 | 2.84±0.21 | 5.44±0.29 | 2.82±0.19 | 5.68±0.88 | 2.49±0.18 | 5.57±0.34 |
| CP-101,606 infusion: high | | 2.55±0.20 | 5.03±0.36 | 2.81±0.28 | 5.54±0.45 | 2.83±0.28 | 5.63±0.40 | 2.88±0.20 | 5.80±0.42 | 2.89±0.15 | 5.57±0.21 |

Behavioural data are presented as mean ± SEM. Drug studies are listed in chronological order for each cohort. Ketamine doses are listed in brackets.

**Table S4 –** Data for accuracy for baseline weeks between drug studies from the JBT.

| Accuracy | | | | | | | | | | | |
|---|---|---|---|---|---|---|---|---|---|---|---|
| | | Day 1 | | Day 2 | | Day 3 | | Day 4 | | Day 5 | |
| Drug | Cohort | HT | LT | HT | LT | HT | LT | HT | LT | HT | LT |
| Memantine | 1 | 96.2±0.80 | 79.5±1.40 | 96.1±0.91 | 84.8±1.31 | 97.2±1.00 | 84.8±1.10 | 97.8±0.44 | 83.1±1.83 | 96.4±1.28 | 82.4±1.30 |
| MK-801 | | 95.9±1.00 | 83.4±1.27 | 95.3±1.22 | 85.4±2.06 | 96.9±0.76 | 86.7±1.11 | 97.5±0.80 | 86.0±1.38 | 97.7±0.51 | 88.8±1.08 |
| Lanicemine | | 97.3±0.60 | 86.4±1.82 | 98.8±0.48 | 88.4±1.48 | 98.0±0.56 | 88.9±1.68 | 98.8±0.43 | 87.9±1.61 | 96.8±0.69 | 82.4±1.44 |
| CP-101,606: low | | 95.8±1.49 | 82.3±1.25 | 97.8±0.65 | 87.8±1.69 | 98.5±0.42 | 88.4±1.31 | 98.2±0.62 | 90.1±1.49 | 98.6±0.46 | 89.3±1.34 |
| CP-101,606: high | | 97.6±0.73 | 84.6±1.75 | 98.0±0.72 | 86.8±2.04 | 97.7±0.93 | 90.2±1.40 | 97.9±0.56 | 89.8±1.79 | 98.6±0.48 | 88.9±1.09 |
| Scopolamine | | 97.5±0.57 | 85.6±1.79 | 98.3±86.6 | 86.6±1.70 | 96.3±1.20 | 87.8±1.93 | 98.0±0.57 | 86.3±1.33 | 97.1±0.64 | 87.1±1.22 |
| PCP | 2 | 91.0±3.68 | 79.3±2.05 | 93.8±2.03 | 81.5±2.56 | 97.1±1.48 | 82.1±2.07 | 97.8±1.01 | 81.6±1.87 | 96.6±1.78 | 80.9±2.86 |
| Ketamine (10) | | 97.3±0.94 | 82.4±3.36 | 98.6±0.57 | 85.9±2.46 | 99.5±0.35 | 85.5±1.60 | 98.9±0.58 | 85.0±1.86 | 98.9±0.57 | 86.7±1.12 |
| Ketamine (25) | | 93.5±3.66 | 79.6±3.50 | 98.1±0.88 | 84.5±3.28 | 97.0±1.08 | 86.5±2.18 | 98.7±0.79 | 87.8±1.76 | 97.6±1.19 | 85.1±2.23 |
| Post-Surgery | 3 | 92.2±1.47 | 70.4±1.73 | 94.9±1.11 | 85.4±1.75 | 95.1±1.13 | 84.9±1.45 | 93.2±1.29 | 87.8±1.64 | 93.1±1.68 | 86.6±1.93 |
| Ketamine (1) | | 93.0±2.98 | 79.1±3.42 | 93.6±3.03 | 85.0±2.66 | 94.2±3.06 | 86.1±2.62 | 91.9±2.03 | 82.1±1.22 | 93.0±1.45 | 81.6±1.76 |
| Infusions 1 | | 96.3±1.22 | 81.7±2.50 | 96.7±0.89 | 83.6±2.10 | 96.0±0.80 | 86.6±2.23 | 96.9±0.79 | 88.6±1.78 | 96.7±0.45 | 84.7±1.24 |
| CP-101,606 infusion: low | | 97.3±0.66 | 86.8±1.56 | 94.8±1.41 | 87.1±1.90 | 96.7±1.20 | 85.3±1.92 | 97.9±0.88 | 86.2±2.18 | 95.8±0.98 | 86.1±2.45 |
| CP-101,606 infusion: high | | 96.9±0.79 | 83.6±1.89 | 96.5±0.90 | 82.3±1.81 | 98.0±0.77 | 84.5±0.90 | 97.3±0.74 | 87.0±1.63 | 97.0±0.81 | 85.4±1.58 |

Behavioural data are presented as mean ± SEM. Drug studies are listed in chronological order for each cohort. Ketamine doses are listed in brackets.

**Table S5 –** Data for omissions for baseline weeks between drug studies from the JBT.

| Omissions | | | | | | | | | | | |
|---|---|---|---|---|---|---|---|---|---|---|---|
| | | Day 1 | | Day 2 | | Day 3 | | Day 4 | | Day 5 | |
| Drug | Cohort | HT | LT | HT | LT | HT | LT | HT | LT | HT | LT |
| Memantine | 1 | 0.00±0.00 | 2.57±0.86 | 0.66±0.32 | 2.55±0.82 | 0.26±0.18 | 3.84±0.86 | 0.00±0.00 | 3.86±1.10 | 0.42±0.30 | 4.32±0.96 |
| MK-801 | | 1.36±0.70 | 3.91±0.95 | 0.84±0.41 | 6.30±1.60 | 0.93±0.43 | 3.30±1.16 | 0.68±0.43 | 2.64±0.93 | 0.32±0.22 | 3.93±1.18 |
| Lanicemine | | 0.38±0.28 | 1.98±0.52 | 0.00±0.00 | 3.13±1.08 | 0.13±0.13 | 2.44±0.82 | 0.28±0.19 | 3.94±1.17 | 0.27±0.18 | 4.34±0.97 |
| CP-101,606: low | | 0.87±0.44 | 3.61±1.02 | 0.53±0.24 | 3.18±1.03 | 0.27±0.27 | 2.50±0.73 | 0.38±0.27 | 1.07±0.34 | 0.14±0.14 | 3.98±1.05 |
| CP-101,606: high | | 0.77±0.37 | 5.77±1.53 | 0.26±0.17 | 4.48±1.00 | 0.24±0.24 | 4.03±1.14 | 0.52±0.29 | 3.67±1.13 | 0.26±0.18 | 3.52±0.90 |
| Scopolamine | | 0.40±0.29 | 3.10±0.94 | 0.00±0.00 | 3.55±0.95 | 0.26±0.18 | 3.90±0.75 | 0.13±0.13 | 5.07±1.26 | 0.89±0.45 | 5.77±1.16 |
| PCP | 2 | 0.97±0.64 | 3.32±1.35 | 0.87±0.58 | 8.05±2.58 | 0.25±0.25 | 6.51±2.19 | 0.54±0.36 | 5.07±2.04 | 0.84±0.51 | 5.87±1.18 |
| Ketamine (10) | | 0.00±0.00 | 5.28±1.91 | 0.54±0.36 | 7.75±1.90 | 0.00±0.00 | 5.35±0.85 | 0.28±0.28 | 6.62±2.37 | 0.00±0.00 | 3.49±0.89 |
| Ketamine (25) | | 0.85±0.59 | 6.00±2.17 | 0.00±0.00 | 5.71±1.89 | 0.81±0.57 | 4.68±1.46 | 0.26±0.26 | 4.89±1.66 | 0.27±0.27 | 8.32±2.21 |
| Post-Surgery | 3 | 0.91±0.40 | 3.50±0.99 | 0.53±0.44 | 3.28±0.89 | 0.64±0.42 | 2.25±0.41 | 0.52±0.23 | 2.87±0.68 | 0.42±0.23 | 2.76±0.77 |
| Ketamine (1) | | 0.00±0.00 | 5.95±1.94 | 0.14±0.14 | 2.03±0.64 | 0.07±0.07 | 2.17±0.63 | 0.13±0.13 | 3.32±0.68 | 0.27±0.19 | 2.33±0.55 |
| Infusions 1 | | 0.00±0.00 | 3.96±1.43 | 0.35±0.24 | 3.17±1.42 | 0.19±0.19 | 5.58±1.46 | 0.00±0.00 | 3.24±1.24 | 0.14±0.14 | 4.12±1.85 |
| CP-101,606 infusion: low | | 0.16±0.16 | 4.14±0.90 | 0.48±0.25 | 3.53±0.95 | 0.54±0.37 | 3.94±1.07 | 0.16±0.16 | 5.07±1.26 | 0.00±0.00 | 5.01±1.01 |
| CP-101,606 infusion: high | | 0.83±0.67 | 4.82±1.41 | 0.33±0.22 | 4.01±1.16 | 0.65±0.37 | 4.86±1.15 | 0.16±0.16 | 4.29±1.28 | 0.54±0.37 | 4.25±0.98 |

Behavioural data are presented as mean ± SEM. Drug studies are listed in chronological order for each cohort. Ketamine doses are listed in brackets.

**Table S6** – Data for premature responding for baseline weeks between drug studies from the JBT.

| Premature | | | | | | |
|---|---|---|---|---|---|---|
| **Drug** | **Cohort** | **Day 1** | **Day 2** | **Day 3** | **Day 4** | **Day 5** |
| Memantine | 1 | 7.71±1.56 | 6.38±0.97 | 6.19±0.90 | 7.02±0.83 | 6.30±0.73 |
| MK-801 | | 8.03±1.00 | 6.57±0.69 | 5.77±1.13 | 5.76±0.84 | 5.56±1.01 |
| Lanicemine | | 6.38±0.81 | 5.60±0.83 | 5.57±0.93 | 5.03±0.71 | 8.72±1.37 |
| CP-101,606: low | | 6.58±1.06 | 5.13±0.91 | 4.21±0.48 | 6.07±0.95 | 4.00±0.69 |
| CP-101,606: high | | 5.05±0.63 | 4.91±0.77 | 3.62±0.79 | 4.26±0.560 | 3.41±0.46 |
| Scopolamine | | 5.81±1.15 | 4.71±0.83 | 4.05±0.72 | 3.26±0.62 | 4.80±0.91 |
| PCP | 2 | 5.81±1.20 | 4.39±0.95 | 2.73±0.94 | 3.86±1.13 | 4.17±0.49 |
| Ketamine (10) | | 12.0±2.15 | 6.55±1.62 | 3.66±0.96 | 3.13±0.49 | 3.44±0.88 |
| Ketamine (25) | | 11.76±1.39 | 6.20±1.24 | 5.25±0.98 | 6.45±1.09 | 6.84±1.73 |
| Post-Surgery | 3 | 39.4±3.86 | 19.9±2.16 | 14.1±1.88 | 13.8±1.98 | 13.0±1.89 |
| Ketamine (1) | | 14.0±4.41 | 11.8±3.03 | 12.3±3.00 | 13.0±1.89 | 12.3±2.00 |
| Infusions 1 | | 11.2±1.98 | 6.27±0.74 | 6.46±1.11 | 7.26±1.25 | 6.58±1.64 |
| CP-101,606 infusion: low | | 6.74±0.87 | 6.39±1.03 | 4.86±0.82 | 4.89±0.80 | 5.86±0.98 |
| CP-101,606 infusion: high | | 5.31±0.76 | 3.85±0.72 | 5.38±0.78 | 4.98±1.00 | 4.35±0.75 |

Behavioural data are presented as mean ± SEM. Drug studies are listed in chronological order for each cohort. Ketamine doses are listed in brackets.

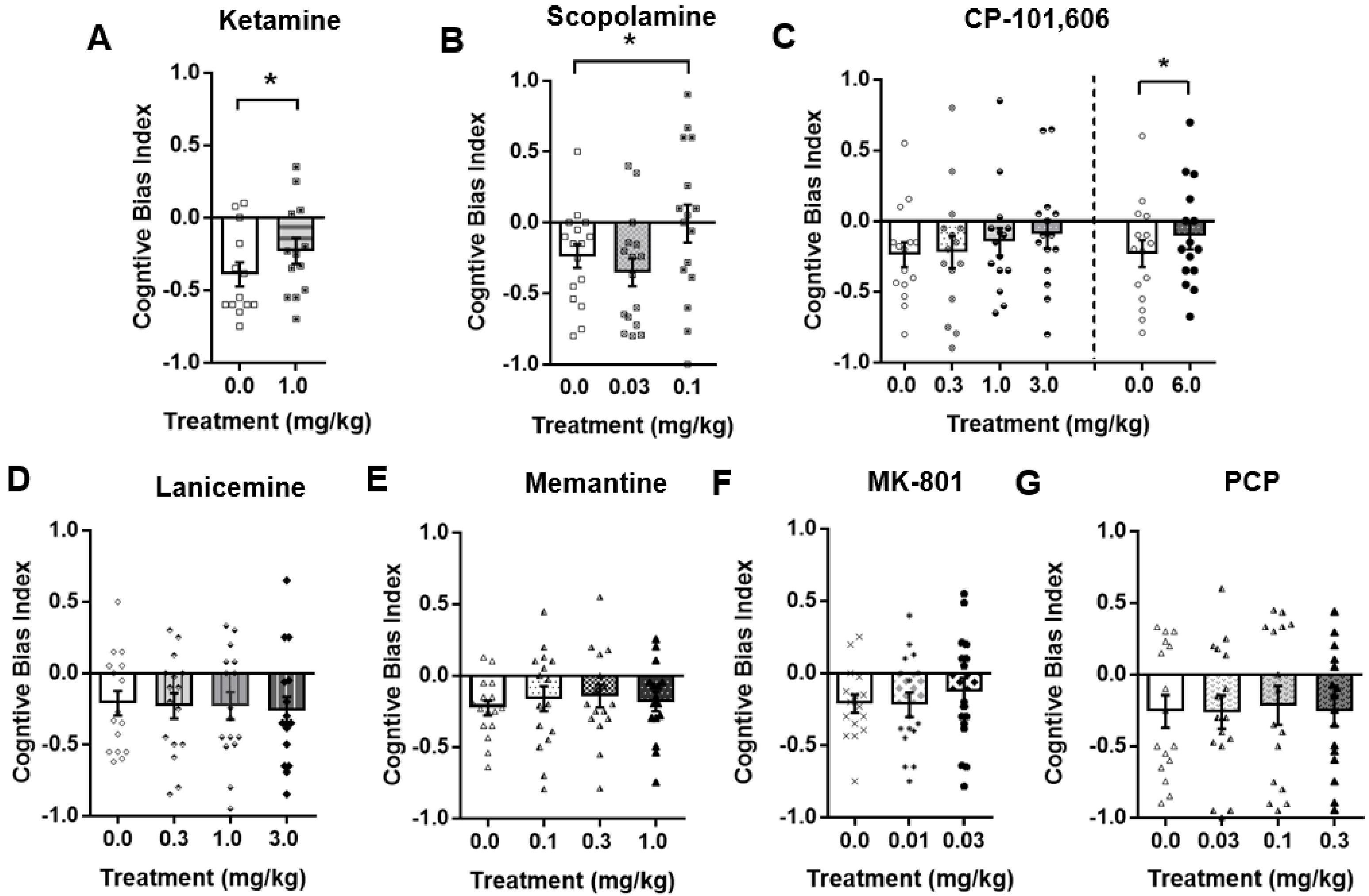


**Figure S2** – *The effect of acute treatment with rapid acting antidepressant drugs and NMDA receptor antagonists on judgement bias of the midpoint ambiguous tone displayed as CBI.*

This figure shows data from Figure 1 displayed as cognitive bias index (CBI) scores. Ketamine (0.0, 1.0 mg/kg; n = 13), scopolamine (0.0, 0.03, 0.1 mg/kg; n = 16), CP-101,606 (Expt 1: 0.0, 0.3, 1.0, 3.0 mg/kg, n = 15; Expt 2: 0.0, 6.0 mg/kg, n = 15), lanicemine (0.0, 0.3, 1.0, 3.0 mg/kg; n = 16), memantine (0.0, 0.1, 0.3, 1.0 mg/kg; n = 16) and MK-801 (0.0, 0.01, 0.03 mg/kg; n = 16 were administered acutely by intraperitoneal injection prior to testing on the judgement bias task. (A) Replicating previous studies, ketamine (1.0 mg/kg) caused CBI to shift in the positive direction. (B) Scopolamine (0.1 mg/kg) changed CBI scores to near zero, a positive shift. (C) CBI became moved in a positive direction after a 6.0 mg/kg dose of CP-101,606. (D-G) Lanicemine, memantine, MK-801 and low doses of PCP did not alter CBI for the midpoint tone at the doses tested. Data shown and represent mean ± SEM (bars and error bars) overlaid with individual data points for each rat. Dashed line (panel C) indicates separate, counterbalanced experiments. *$p$ < 0.05. CP-101,606, ketamine, lanicemine, memantine, PCP: 60 min pre-treatment; scopolamine, MK-801: 30 min pre-treatment.

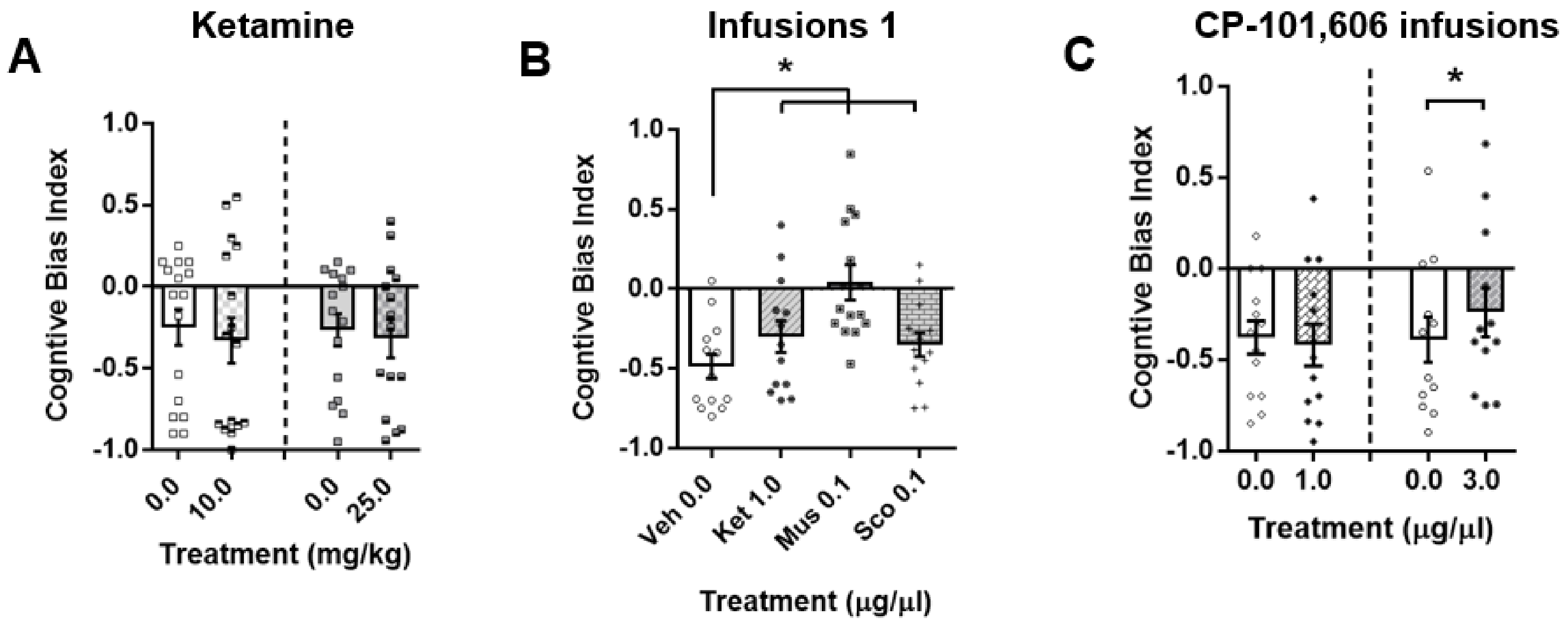


**Figure S3** – *The effect of acute treatment with high doses of ketamine, and mPFC infusions of rapid acting antidepressants on judgement bias of the midpoint ambiguous tone displayed as CBI.*

This figure shows data from Figures 2-4 displayed as cognitive bias index (CBI) scores. (A) From Figure 2: ketamine (Expt 1: 0.0, 10.0 mg/kg, n = 16; Expt 2: 0.0, 25.0 mg/kg, n = 16) were administered by intraperitoneal injection to measure their effect on judgement bias. Neither dose had any effect on CBI. (B) From Figure 3: ) In the first infusion experiment, ketamine (Ket; 1.0 µg/µl) muscimol (Mus; 0.1 µg/µl), scopolamine (Sco; 0.1 µg/µl) or vehicle (Veh; 0.0 µg/µl; n = 13), were administered by intracerebral infusion into the mPFC to measure the effect on judgement bias. All infuions caused positive changes in CBI. (C) From Figure 4: CP-101,606 (Expt 1: 00.0, 1.0 µg/µl, n = 13; Expt 2: 0.0, 3.0 µg/µl, n = 12) was administered by intracerebral infusion in the mPFC to measure the effect on judgement bias. Only the higher dose (3.0 mg/kg) caused CBI to become more positive. Data shown and represent mean ± SEM (bars and error bars) overlaid with individual data points for each rat. Dashed line (panel C) indicates separate, counterbalanced experiments. *$p < 0.05$. Ketamine (systemic): 60 min pre-treatment; infusions: 5 min pre-treatment.

**Table S7** – Description of statistical analysis for other behavioural measures

| Behavioural measure | Analysed for: | Description | Statistical analysis |
|---|---|---|---|
| **Response latency** | Each tone | Time between presentation of the tone and response on the lever (correct lever for high and low reward tones, either lever for midpoint tone) | Two-way repeated measures ANOVA with tone and session as within-subjects factors |
| **Accuracy** | Reference tones (high and low tones) | Number of correct responses made divided by the total number of responses made (correct + incorrect) for that tone | |
| **Percentage omissions** | Each tone | Number of trials where no lever press occurred during 20 s tone presentation divided by total completed trials for that tone | |
| **Percentage of premature responses** | Whole session | Number of trials where a response was made in the 5 s inter-trial interval divided by total completed trials | Repeated measures ANOVA with session as the within-subjects factor |

This table details the other behavioural measures (apart from cognitive bias index) that were analysed for each experimental manipulation and are presented in Table 1. ANOVA – analysis of variance.

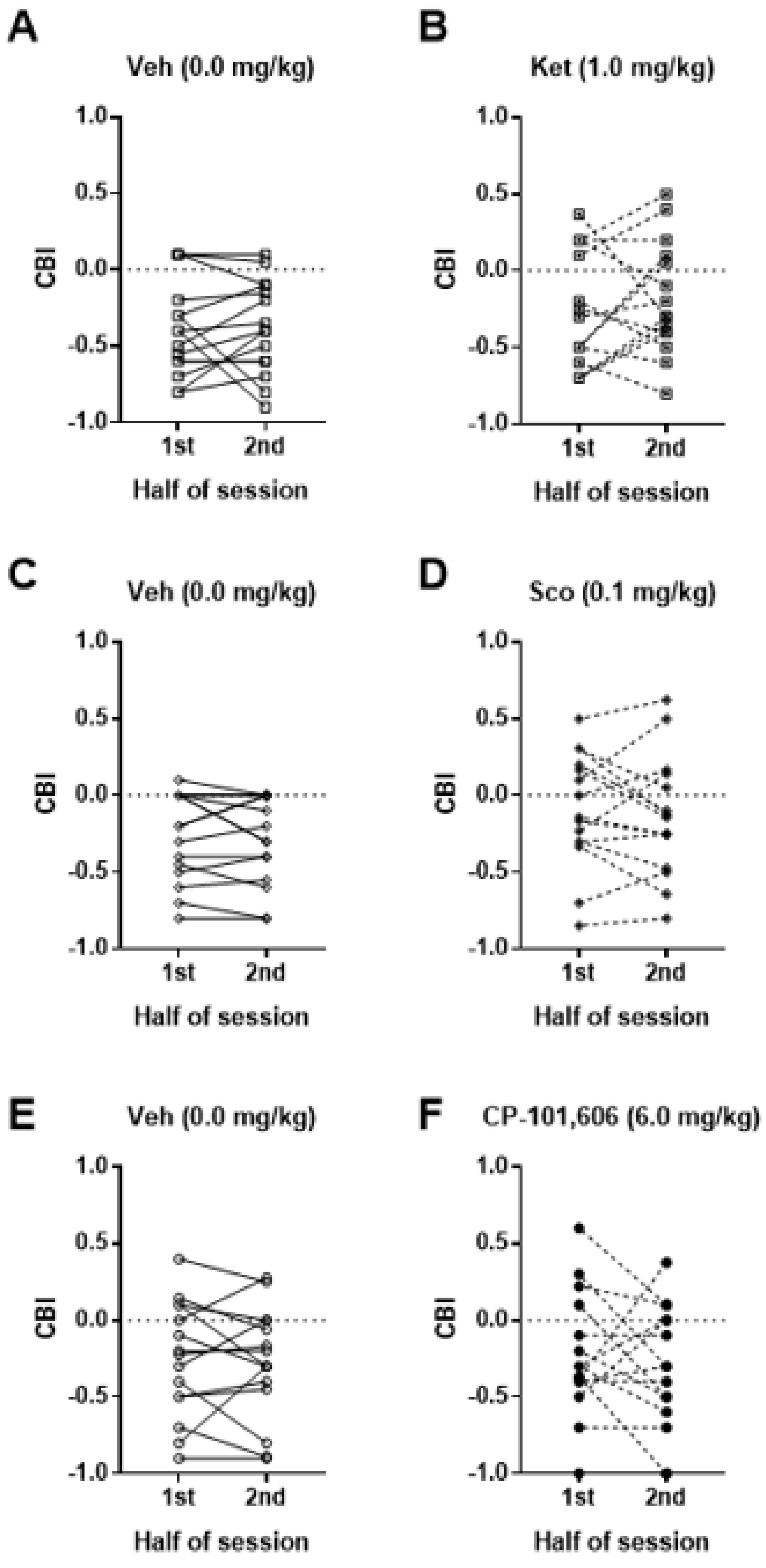


**Figure S4 – CBI analysed by session split in half for systemic drugs that caused a positive change in bias.**

Ketamine (0.0, 1.0 mg/kg; n = 13), scopolamine (0.0, 0.03, 0.1 mg/kg; n = 16), CP-101,606 (Expt 2: 0.0, 6.0 mg/kg, n = 15) were administered acutely by intraperitoneal injection prior to testing on the judgement bias task. Data from these drug studies for doses that showed a positive change in judgement bias (ketamine: 1.0 mg/kg; scopolamine: 0.1 mg/kg) and CP-101,606: 6.0 mg/kg) were re-analysed by splitting each session in half, and comparing CBI for the first and last half of the sessions. Vehicle doses for each drug (panels A,C,E) are also shown for comparison. There is no consistent change between CBI scores across the first and second halves of a session for the drugs shown. Data shown are individual data points, linked for each individual rat.
